# Pharmacokinetic/pharmacodynamic analysis of oral amoxicillin-clavulanic acid against extended-spectrum beta-lactamase producing *Escherichia coli* in an *in vitro* bladder infection model: impact on clinical breakpoints

**DOI:** 10.64898/2026.11.12.766335

**Authors:** Connor RB Anderson, Elke van Gorp, Hayoung Won, Jason A Roberts, Hugh Cottingham, Siobhonne KJ Breen, Cornelia B Landersdorfer, Joseph Meletiadis, Anton Y Peleg, Iain J Abbott

**Author notes:** Corresponding author: Address correspondence to Dr Iain J Abbott, Qualifications: MBBS PhD FRACP FRCPA, Full address: The Alfred, 55 Commercial Road, Melbourne VIC 3004.

## Abstract

Uropathogens with acquired ESBL-genes can limit effective oral treatment options. Amoxicillin with clavulanic acid can be used as targeted therapy (500/125mg, 8-hourly). Despite favorable urinary pharmacokinetics (PK), there is little urine-specific data that quantifies the exposure required for bacterial clearance. A dynamic *in vitro* bladder model was employed, allowing for the simulated treatment of UTIs with oral amoxicillin-clavulanic acid. Sixteen *Escherichia coli* isolates (12 clinical ceftriaxone-resistant uropathogens, 4 laboratory isolates) were selected. Bacterial response to treatment, emergence of resistance, exposure-response targets and PK/PD indexes were explored. Following simulation of standard-dose oral amoxicillin-clavulanic acid (500/125mg, 8-hourly; C_max_ 935mg/L; AUC_0-24_ 14272mg/L), 14/16 isolates displayed no growth at the end of treatment, with elimination in seven isolates confirmed after a 24h antibiotic-free follow up. As exposure was reduced, greater survival and regrowth were observed. Only two isolates had significant post-exposure rises in MIC (32 to 128mg/L, 64 to 256mg/L). %T_>MIC_ was the best predictor for end of treatment outcome (Δlog_10_cfu/mL: stasis 52.7%, 1log_10_ kill 61.8%, 2log_10_ kill 71.1%, R^2^=0.9062). Lower R^2^ values were found with EUCAST MIC measurements (fixed 2mg/L clavulanic acid) as compared to CLSI MIC measurements (2:1 amoxicillin to clavulanic acid). Monte Carlo simulations indicated that standard-dose 8-hourly would achieve all %T_>MIC_ targets in >95% of virtual patients with MICs ≤64 mg/L, regardless of MIC method (CLSI or EUCAST). Amoxicillin-clavulanic acid displays time-dependent killing against ceftriaxone-resistant *E. coli*. This data supports clinical breakpoints for uncomplicated UTI for isolates with MIC ≤ 64mg/L, assuming 500/125mg oral dosing 8-hourly.

## INTRODUCTION

Urinary tract infections (UTIs) are responsible for 400-million cases a year globally (1), placing a significant burden on health systems (2). More than 50% of women will experience at least one UTI in their lifetime, with 10% experiencing at least one infection annually (3, 4). The increasing aging population is at greater risk (5), with approximately 30% of women aged ≥ 85 years old experiencing at least one UTI in the previous year (6). Similarly, 35-40% of kidney transplant patients experience a UTI following transplant (7–9). UTIs represent 1.2% of all conditions managed by general practices in Australia (10), with empiric antibiotics prescribed in 90% of cases (11). Whilst antibiotic resistance rates remain low to first line agents such as nitrofurantoin and fosfomycin, resistance rates to trimethoprim and ciprofloxacin are increasing (11), and the acquisition of extended-spectrum beta-lactamases (ESBL) enzymes, particularly in *Escherichia coli* ST131, limit treatment options (12, 13). *E. coli* causes 65-80% of uncomplicated UTIs (11, 14, 15), with ESBL-producing *E. coli* associated with bloodstream infections, and a with a higher rates of patient mortality (OR = 1.70) (16) and longer length of stay in hospital (17, 18). Compounding the issue, ESBL-producers have high rates of co-resistance to other antibiotics, limiting treatment options significantly (19).

Amoxicillin is a beta-lactam antibiotic, which can be co-formulated with the beta-lactamase inhibitor clavulanic acid. The addition of clavulanic acid can restore the bactericidal activity of amoxicillin against some ESBL-producing organisms, such as TEM-, SHV- and CTX-M-type ESBLs (20). However, co-production of clavulanic acid-resistant enzymes (particularly OXA-1), β-lactamase hyperproduction and porin loss can mean ESBL-producing isolates can test non-susceptible. Inhibition is also dependent on clavulanic acid concentration and the ratio of amoxicillin to clavulanic acid at the site of infection (21). Despite high oral bioavailability, ability to concentrate in the urine, and antibacterial activity largely unaffected by pH, amoxicillin- clavulanic acid has remained a second line or reserve therapy for uncomplicated UTIs (21, 22). This is primarily driven by concerns of its lower efficacy compared to first-line agents (nitrofurantoin, fosfomycin) and antimicrobial stewardship principles limiting the use of broad-spectrum agents. Uncomplicated UTIs clinical and microbiological cure rates for amoxicillin and amoxicillin-clavulanic acid have been found to be as low as 58% and 67-88% respectively (23, 24). These studies have been widely cited as evidence against the use of amoxicillin-clavulanic acid, however, the dosing regimens in these studies were conducted for short durations (3-days) with infrequent dosing intervals (BID). Pharmacokinetic/pharmacodynamic (PK/PD) evidence demonstrates that beta-lactam antibiotics display time-dependent bactericidal activity, supporting more frequent dosing (25, 26). Despite concerns about the clinical efficacy of beta-lactam/beta lactamase inhibitor combinations for ESBL-producing bacteria, there have been renewed efforts to utilize amoxicillin-clavulanic acid for antibiotic resistant UTIs. ESBL-producing organisms often still show *in vitro* susceptibility to the amoxicillin-clavulanic acid combination. A UK-based study found 66.3% of ESBL-producing *Enterobacterales* displayed amoxicillin-clavulanic acid susceptibility (27). In a Swedish based study, CTX-M group 1 and CTX-M group 9 organisms displayed amoxicillin-clavulanic acid susceptibility rates upwards of 80% and 95% respectively (28).

Patient factors, drug pharmacokinetics, and the co-acquisition of resistance to multiple classes of antibiotics can greatly limit viable oral treatment options. The IDSA 2026 Guidance on the treatment of Antimicrobial-Resistant Gram-Negative Infections does not suggest the use of amoxicillin-clavulanic acid for uncomplicated UTIs caused by ESBL-producing *Enterobacterales* (29), stating that although clavulanic acid inhibits ESBL enzymes *in vitro*, this may not translate into clinical efficacy. To address the lack of modern urinary PK/PD data supporting the amoxicillin-clavulanic acid combination for this key pathogen group, we have performed PD profiling of oral amoxicillin-clavulanic acid employing a dynamic bladder infection *in vitro* model. We investigated the PK/PD indices for amoxicillin-clavulanic acid activity against ceftriaxone-resistant *E. coli* urinary isolates to inform UTI-specific clinical breakpoints.

## MATERIALS & METHODS

### Media and antibiotics

Cation-adjusted Mueller-Hinton agar (CAMHA; Becton-Dickinson, USA) was used for bacterial quantification and disk diffusion susceptibility testing. Cation-adjusted Muller-Hinton Broth (CAMHB; Becton-Dickinson, USA), pooled human urine (collected with ethical approval; project #27033), and modified synthetic human urine (mSHU; pH 5.6; table S1) (30) were used as liquid media for broth microdilution (BMD) susceptibility testing and bacterial growth experiments. Antibiotic disks (10 μg ampicillin and 20/10 μg amoxicillin-clavulanic acid; Oxoid Ltd/Thermo Fisher Scientific, UK) were utilized for disk diffusion susceptibility testing. Amoxicillin (Merck Germany, A8523), potassium clavulanic acid (Merck Germany, 3345) and lithium clavulanic acid (Merck Germany, L0720000) powders were utilized for broth microdilution (BMD) susceptibility testing and to produce amoxicillin-clavulanic acid-supplemented agar plates (AMC-CAMHA-32/2 mg/L). Parental formulations of amoxicillin-clavulanic acid (2000mg-200mg; Curam, Sandoz, Switzerland) and amoxicillin (1000mg; Ibiamox, Juno Pharmaceuticals, Australia) were used to generate antibiotic exposures within the bladder infection model. Media and antibiotic stock generation methodologies are described in the supplementary methods.

### *E. coli* isolates and susceptibility testing

Ceftriaxone-resistant clinical *E. coli* urinary isolates from a UTI-biorepository (ethical approval; #104833) underwent amoxicillin-clavulanic acid MIC testing by BMD using EUCAST methodology. Ceftriaxone resistance (MIC ≥64mg/L) was used as a surrogate marker for ESBL-production, which was pre-determined by Vitek2 (bioMérieux, France; AST-N344 card). Sixteen isolates were selected, including: 12 ceftriaxone-resistant clinical isolates representing a broad range of baseline amoxicillin-clavulanic acid susceptibility, 2 laboratory QC strains (ATCC 25922 and ATCC 35218), and 2 strains used in murine UTI models (CFT073 and UTI89). These isolates underwent further BMD MIC testing in triplicate. Both EUCAST (clavulanic acid concentration fixed at 2 mg/L) and CLSI (2:1 ratio of amoxicillin-clavulanic acid) BMD methodologies and interpretations were employed (31, 32). Additional BMD susceptibility testing was performed in nonstandard media (mSHU and pooled human urine). Antimicrobial disk diffusion susceptibility testing was performed in duplicate (average inhibition zone presented). Whole genome sequencing (WGS) was performed at baseline on the sixteen isolates. Achtman sequence type (ST) and beta-lactamase genes were identified (see supplementary methods).

### Bladder infection model set up

The bladder infection model (BIM) is a dynamic, multi-compartmental, open dilutional *in vitro* system adapted from Abbott *et al.* (33) (Figure 1). It simulates urodynamics (constant bladder fill with intermittent voiding) and urinary antibiotic exposure following humanized dosing schedules. This enabled the simulation of uncomplicated UTI treatment with bacteria held within a urinary matrix. Total flow rate of mSHU liquid medium was set to 400 mL/h. The volume of the intestinal and circulatory compartments was maintained at 350 mL and 1000 mL, respectively. Sixteen bladder compartments were run concurrently, with an individual in-flow rate of 25 mL/h, with a 4-hourly voiding schedule. The BIM is described in greater detail in the supplementary methods.

**Figure 1.**
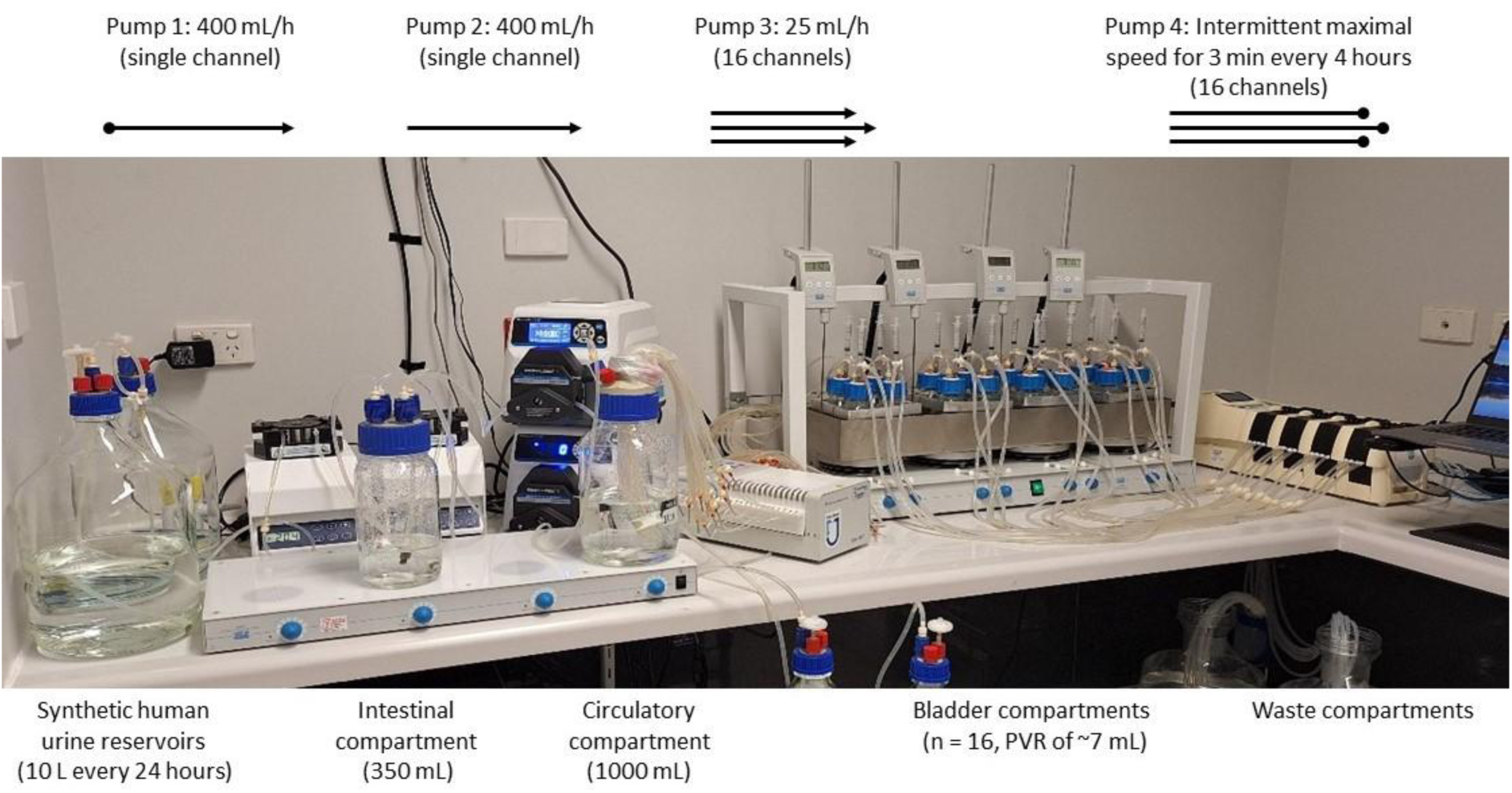
The bladder infection model (BIM). A dynamic, multicompartmental, dilutional *in vitro* model used for the simulation of an uncomplicated urinary tract infection and treatment with amoxicillin-clavulanic acid following oral dosing. Synthetic human urine flows from fresh media reservoirs (left; 10 L of media used per day) to compartment A (‘intestinal’ compartment; into which antibiotics are administered) to compartment B (‘circulatory’ compartment), then split to deliver media with dynamically changing antibiotic concentrations into 16 individual bladder compartments, held in a water bath at 37°C. The volumes of the intestinal and circulatory compartments are kept static, providing tandem first-order processes to simulate antimicrobial absorption and excretion. The media volume in each bladder compartment is allowed to increase over time at 25 mL/h, until a timed voiding program empties each bladder down to a post void residual volume (∼7 mL). A 24 h antibiotic reservoir is prepared daily (held at 4-8°C), with timed dosing into compartment A. Prior to each repeat dose of antibiotic, the contents within compartment A are automatically voided. The model runs continuously for 144 hours.

### Simulated amoxicillin-clavulanic acid exposure

Human urinary amoxicillin concentrations observed after oral dosing from the published literature were utilized to inform the target urinary exposure within the BIM (C_max_, C_min_, AUC_0-24_) (34). The full-dose simulation in the BIM considered average values within the wide interindividual variation observed in human urinary amoxicillin exposures (peak urinary concentrations are 306-856 mg/L after 250 mg oral amoxicillin, and 115-1,850 mg/L after 500 mg oral amoxicillin; 80% bioavailability; 60% of the dose excreted unchanged in urine in the first 6 h) (34). Clavulanic urinary exposure is not well described in the literature, with highly variable absorption, 50% hepatic clearance, 30% protein binding, and only 28% of the dose excreted in urine by 6-hours (35, 36). Without clear urinary targets for clavulanic acid, and for practical reasons, amoxicillin-clavulanic acid was administered using the parental formulation with a 10:1 ratio of amoxicillin to clavulanic acid. Drug distribution equations were used to calculate the required media flow rates (mL/h), compartment volumes (mL) and dose of amoxicillin-clavulanic acid (mg) (37).

Dosing simulated a 5-day treatment course. To quantify an exposure-response relationship, experiments were performed using a stepwise reduction in exposure. The initial, full-dose experiment simulated an 8-hourly antibiotic dosing to generate (at steady state) a urinary amoxicillin peak concentration (C_max_) of 935 mg/L, and area-under-the curve (AUC_0-24_) of 14272 mg/L, and 86% of the dose excreted over the first 6-hours (equivalent to humans when bioavailability and the absorption phase is considered). Subsequent BIM runs targeted amoxicillin C_max_ of 467 mg/L and 220 mg/L, with a frequency of dosing of 8-hourly, 12-hourly and daily. An additional ‘amoxicillin-only’ experiment targeted a C_max_ 220 mg/L with 8-hourly dosing, to enable direct comparison with outcomes following the same amoxicillin exposure with clavulanic acid.

### Antibiotic quantification

Amoxicillin and clavulanic acid concentrations were measured by a validated ultra-high-performance liquid chromatography-tandem mass spectrometry (UHPLC-MS/MS) method on a Nexera2 liquid chromatograph connected to an 8030+ triple quadrupole mass spectrometer (Shimadzu, Kyoto, Japan). Test samples were passed through a 0.22-micron PES filter (MilliporeSigma, United States, SLGPR33RS) prior to immediate storage at -80°C. Samples were then tested in batches alongside calibrators and quality controls, with results subject to batch acceptance criteria (US FDA, M10 Bioanalytical Method Validation and Study Sample Analysis, Guidance for Industry, November 2022) (38). The calibration range was 0.1 to 200 mg/L for amoxicillin, and 0.1 to 20 mg/L for clavulanic acid. Samples that exceeded the upper limit of quantification were pre-diluted with mSHU. Assessment of accuracy and precision was performed to meet the US FDA guidelines for bioanalytical method validation. Supplementary HPLC-PDA was performed to analyze peak areas for amoxicillin and amoxicillin-degradation products. Stability of both antibiotics over 24 h was assessed by suspending 1000mg of amoxicillin and 100mg of clavulanic acid in mSHU at 4-8°C, room temperature, and 37°C.

### Drug-free static and dynamic growth kinetics

Static growth assays were performed under drug free conditions to assess bacterial growth rate and total growth capacity of the isolates in different media. Four isolates, ATCC 25922, ATCC 35218, CFT073 and UTI89 were inoculated at 10^5^ CFU/mL and grown in 20 mL of CAMHB, pooled human urine, and mSHU media at 37°C. Following this, the BIM was run in mSHU for 24 hours without antibiotics to determine the drug-free maximal growth capacity of all 16 isolates under dynamic conditions.

### Bacterial response in the BIM

Before the first amoxicillin-clavulanic acid dose, the BIM was run for 2 hours to allow bacteria to enter an exponential growth phase before antibiotic exposure. Following antibiotic exposure, an end of treatment (EOT) assessment was performed at 120 h. The model was continued for an additional 24 h (up to 144 h) without further drug exposure, to determine if complete bacterial elimination was achieved at follow-up (F/U). Quantitative cultures on CAMHA were used to determine bacterial density (cfu/mL) in each bladder compartment, sampled prior to antibiotics, twice a day during therapy and the follow-up period. The limit of detection was 50 cfu/mL. At baseline, 120 h and 144 h, samples were additionally plated onto antibiotic-supplemented agar (AMC-CAMHA-32/2 mg/L) to quantify emergence of resistant subpopulations (see Supplementary Methods). Post-exposure amoxicillin-clavulanic acid BMD MIC susceptibility testing was performed where there was bacterial growth at 120 and 144 h, from both the drug-free and antibiotic-supplemented agar. Emergence of resistance was determined to be significant if the MIC rise from baseline was ≥ 2-fold doubling dilutions.

### PK/PD analysis

The primary PD endpoint was the change in bacterial density at EOT (120 h Δ log_10_ cfu/mL). The secondary PD endpoint was the area-under-the-bacterial-kill-curve at follow-up (AUBKC_0–144_). The PK/PD indices evaluated were AUC_0–24_/MIC and %T _> MIC_. The analysis was performed using both CLSI and EUCAST MIC values. The exposure-response relationship was assessed via a four parameter E_max_ non-linear regression model. This analysis used the PD outcome data from all amoxicillin-clavulanic acid exposure experiments with the simulated PK exposures determined by the mathematical model. R^2^ values were reported to describe the goodness-of-fit. Exposures required for stasis, 1, 2 and 3 log_10_ kill (for Δ log_10_ cfu/mL) as well as stasis, 1, 2 and 3 log_10_ reduction in total bacterial response (AUBKC_0–144_), and the IC_50_ values, were determined. No constraints were applied to the E_max_ non-linear regression model. The amoxicillin only experiment was not included in PK/PD analysis. All data was analyzed, and figures generated using GraphPad Prism (version 11.0.1).

### Monte Carlo simulations (MCS)

MCS were conducted to estimate the probability of target attainment (PTA) across the *E. coli* amoxicillin-clavulanic acid MIC distribution range. Berkeley Madonna software (version 8.3.23) was used to simulate 5,000 virtual patients. The simulated *in vitro* PK profiles for standard (500 mg, 8-hourly), intermediate (500 mg, 12-hourly) and low dose (250 mg, 12-hourly) amoxicillin exposure, with ±50% allowance for variability in human urinary concentrations, were utilized. Clavulanic acid exposure was assumed to be one-tenth of the amoxicillin exposure. Target %T_>MIC_ values associated with bacterial stasis and log killing at EOT were selected in accordance with the PK/PD analysis described in this study. All data were analyzed using GraphPad Prism (version 11.0.1).

## RESULTS

### Bacterial isolates and baseline *in vitro* susceptibility

Eighty-four ceftriaxone-resistant clinical *E. coli* uropathogens underwent amoxicillin-clavulanic acid susceptibility testing using EUCAST methodology. Isolates displayed MICs ranging from 2/2 mg/L to greater than 256/2 mg/L, displaying a bimodal distribution, with peaks at 4 mg/L and 16 mg/L, respectively (Figure 2). The majority (52.4%) of isolates were wild type strains, with MICs below or equal to the epidemiological cut-off value of 8 mg/L. Sixteen isolates were selected and underwent additional BMD MIC susceptibility testing in CAMHB, mSHU and pooled human urine (Table 1). Compared with standard methodology in CAMHB, MIC measurements were lower when tested in urine and mSHU (Figure S1). BMD susceptibility testing for amoxicillin and antimicrobial disk diffusion susceptibility testing was additionally performed, listed in Table S2 & S3.

**Figure 2.**
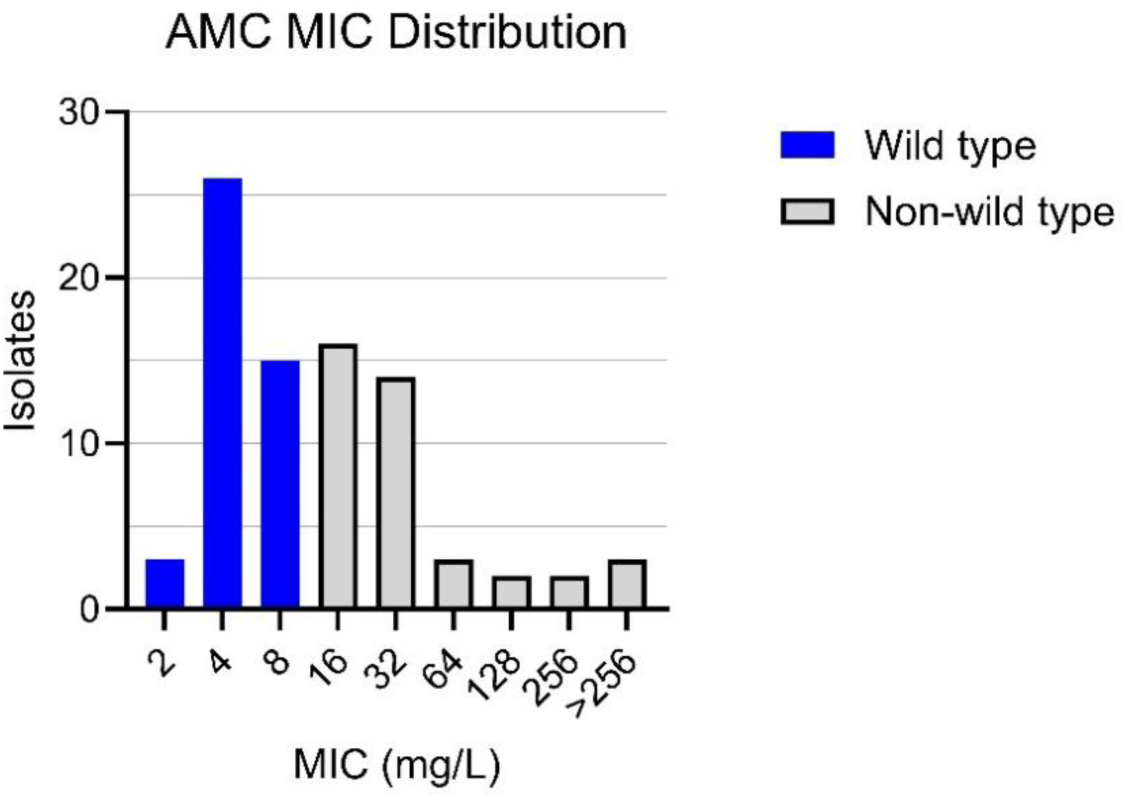
Amoxicillin-clavulanic acid MIC distribution in ceftriaxone-resistant *E. coli* uropathogens. AMC, Amoxicillin-clavulanic acid. AMC MICs of 84 ceftriaxone-resistant *E. coli* clinical uropathogens. Broth microdilution was performed by EUCAST methodology (static 2 mg/L concentration of clavulanic acid). Wild type and non-wild type isolates indicated (epidemiological cut off value, ECOFF is 8 mg/L).

**Table 1.** Amoxicillin-clavulanic acid MICs of sixteen *E. coli* tested in the bladder infection model.

| Isolate no. | ST | β-lactamase gene | Baseline AMC BMD MIC (median (range), mg/L) |  |  |  |  |  |
|---|---|---|---|---|---|---|---|---|
|  |  |  | CAMHB |  | mSHU |  | Human urine |  |
|  |  |  | CLA 2 mg/L | CLA 2:1 | CLA 2 mg/L | CLA 2:1 | CLA 2 mg/L | CLA 2:1 |
| ECO 017 | 219 | CTX-M-15 | 2 (2) | 4 (2 - 4) | 2 (1 - 2) | 2 (2) | 1 (1 - 2) | 2 (2) |
| UTI 089 | 95 | - | 4 (2-4) | 4 (4) | 1 (0.5 - 1) | 1 (1) | 2 (2) | 2 (2) |
| ATCC 25922 | 73 | - | 4 (4) | 4 (4 - 8) | 1 (1 - 2) | 1 (0.5 - 1) | 2 (2) | 2 (2) |
| CFT 073 | 73 | - | 4 (4) | 4 (4) | 1 (1 - 2) | 1 (1) | 2 (2) | 2 (2) |
| ATCC 35218 | 127 | TEM-1 | 8 (8) | 8 (8 - 16) | 2 (2 - 8) | 4 (4 - 8) | 8 (8) | 8 (4 - 8) |
| ECO 104 | 95 | CTX-M-55, TEM-1 | 8 (8) | 8 (8) | 8 (8) | 4 (4 - 8) | 8 (8) | 8 (8) |
| ECO 015 | 131 | CTX-M-27, TEM-1 | 8 (8 - 16) | 8 (8) | 4 (4) | 4 (4) | 8 (8) | 8 (4 - 8) |
| ECO 014 | 38 | CTX-M-14 | 16 (16) | 8 (8) | 4 (4) | 4 (4 - 8) | 8 (8 - 16) | 8 (8) |
| ECO 132 | 648 | CTX-M-65, OXA-10 | 16 (8 - 16) | 8 (8 - 16) | 1 (1) | 2 (2) | 8 (4 - 8) | 8 (4 - 8) |
| ECO 016 | 131 | CTX-M-14, TEM-1 | 16 (16) | 8 (8) | 4 (2 - 8) | 8 (8-16) | 16 (8 - 16) | 8 (4 - 8) |
| ECO 136 | 131 | CTX-M-15, OXA-1 | 32 (32) | 16 (16) | 2 (2) | 4 (4) | 16 (8 - 16) | 8 (8) |
| ECO 108 | 393 | CTX-M-27, TEM-1 | 32 (32) | 16 (8 - 16) | 64 (64 - 128) | 16 (16) | 32 (8 - 32) | 16 (16) |
| ECO 125 | 131 | CTX-M-15, OXA-1 | 32 (32 - 64) | 16 (16) | 1 (1) | 4 (4) | 16 (8 - 16) | 8 (8) |
| ECO 139 | 131 | CTX-M-15, OXA-1 | 64 (64) | 16 (16) | 2 (2 - 4) | 4 (4) | 32 (8 - 32) | 16 (8 - 16) |
| ECO 057 | 131 | CTX-M-27, TEM-1 | 128 (128) | 32 (16 - 32) | 128 (128 - 256) | 32 (32) | 128 (128) | 16 (16) |
| ECO 087 | 1193 | CMY-2, CTX-M-15 | 256 (128 - 256) | 64 (64) | 64 (32 - 64) | 16 (16 - 32) | 256 (256) | 32 (32) |

AMC, Amoxicillin-clavulanic acid. AMX, Amoxicillin. CLA, Clavulanic acid. Amoxicillin and amoxicillin-clavulanic acid BMD performed in standard (CAMHB) and non-standard media (mSHU, human urine) employing both EUCAST (2 mg/L static clavulanic acid) and CLSI (2:1 amoxicillin to clavulanic acid ratio) methodology. All BMD testing was done in triplicate, MIC is given as median (range).

### Simulated amoxicillin-clavulanic acid exposure in the BIM

The UHPLC-MS/MS assays for amoxicillin and clavulanic acid in mSHU demonstrated inter-batch precision of 3.6-9.0% and 2.1-10.5%, respectively and inter-batch accuracy of 0.8-5.2% and 1.2-14.0%, respectively, across three quality control concentrations. During antibiotic stability experiments, little degradation was observed for amoxicillin over the 24-hour period regardless of storage condition (>95% remaining; Figure 3). Clavulanic acid stored at 4-8°C remained stable, showing no sign of degradation at eight hours, and by 24 hours, 97.9% of the clavulanic acid remained. In contrast, when held at 20-22°C and 37°C, clavulanic acid degraded quickly. At 8 hours, 93.7% and 87.7% remained, and by 24 hours 86.0% and 69.2% remained, respectively. During the 5-days antibiotic exposure experiments, measured amoxicillin concentrations matched target values (bias 20.4 ± 37.9 mg/L, 3.4 ± 37.8%; linear regression slope 1.009, R^2^ = 0.9803), as did measured clavulanic acid concentrations (bias 6.4 ± 5.8 mg/L, 16.5 ± 37.0%; linear regression slope 1.093, R^2^ = 0.9686; Figures S2-4). There was minimal inter- and intra-compartmental variability for amoxicillin (4.2 ± 1.1%, and 3.2 ± 1.5%, respectively) and clavulanic acid (7.8 ± 4.0% and 6.1% ± 3.3%, respectively).

**Figure 3.**
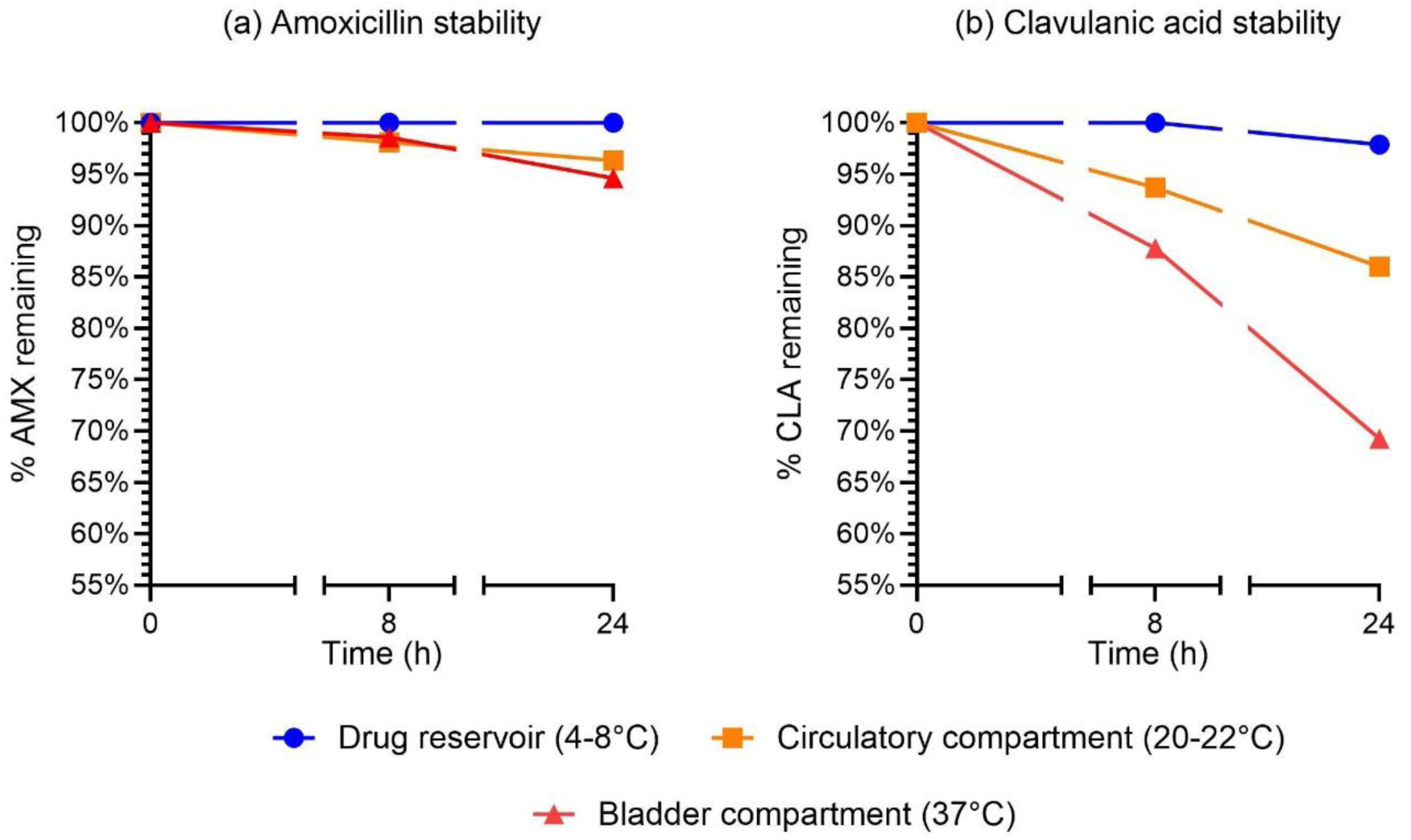
Stability of amoxicillin and clavulanic acid in mSHU. Stability of antibiotics in mSHU under different temperature conditions. Amoxicillin (AMX) remained relatively stable over 24 hours regardless of condition. Clavulanic acid (CLA) stability was dependent on temperature, with colder temperatures resulting in greater stability and warmer temperatures leading to greater instability. Upper limit of the graph has been constrained to 100%.

### Static and dynamic drug-free growth kinetics

Under static drug-free conditions, CAMHB displayed greater growth than in pooled human urine, with faster generation time (t_gen_ 19.2 ± 0.5 min versus t_gen_ 28.5 ± 2.9 min) and higher growth capacity (9.5 ± 0.7 log_10_ CFU/mL versus 8.1 ± 0.1 log_10_ CFU/mL; Figure S5). mSHU and urine displayed comparable static growth kinetics, with a similar generation time (t_gen_ 32.4 ± 3.5 min versus t_gen_ 28.5 ± 2.9 min) and maximal growth capacity (8.2 ± 0.1 log_10_ CFU/mL vs 8.1 ± 0.1 log_10_ CFU/mL). During the dynamic drug free growth experiment in the BIM in mSHU, isolates steadily increased their bacterial density over the 24 h duration despite dilution and bacterial loss because of the urodynamics simulation. Isolates displayed an average maximal drug-free growth capacity in the BIM of 8.0 ± 0.2 log_10_ cfu/mL (Figure S6).

### Exposure-response following amoxicillin-clavulanic acid exposure in the BIM

Following 5 days of full-dose 8-hourly amoxicillin-clavulanic acid simulated therapy, EOT growth was observed in only 2/16 isolates (EUCAST MIC: 128 and 256 mg/L; CLSI MIC: 32/16 and 64/32 µg/mL) (Figure 4; Table 2). After 24 h without additional antibiotic doses, a further 7 isolates displayed growth at F/U. The remaining 7 isolates were eliminated from the bladder compartments, including all isolates with a EUCAST/CLSI MIC ≤ 4 mg/L and 3 isolates with MICs >4 mg/L. Interestingly, ECO139 which was resistant based on EUCAST and intermediate with CLSI was eradicated. Following the half-dose 8-hourly exposure experiment, greater re-growth was observed: 4/16 isolates at EOT (EUCAST MICs ≥8 mg/L, CLSI MIC ≥8/4 mg/L), and 11/16 at F/U. For the half-dose 12-hourly, quarter-dose 12-hourly, and quarter-dose 24-hourly experiments, growth at EOT was observed for 4/16, 11/16 and 16/16 isolates respectively, and all isolates showed regrowth at F/U for all 3 of these exposure experiments. Regrowth was more commonly observed in isolates with an MIC ≥ 8 mg/L. Only two isolates had a significant rise in amoxicillin-clavulanic acid MIC after antibiotic exposure compared to baseline: ECO108 had a rise from 32 to 128 mg/L after half-dose 12-hourly exposure, and ECO139 had a rise of 64 to 256 mg/L after quarter-dose 12-hourly exposure.

**Figure 4.**
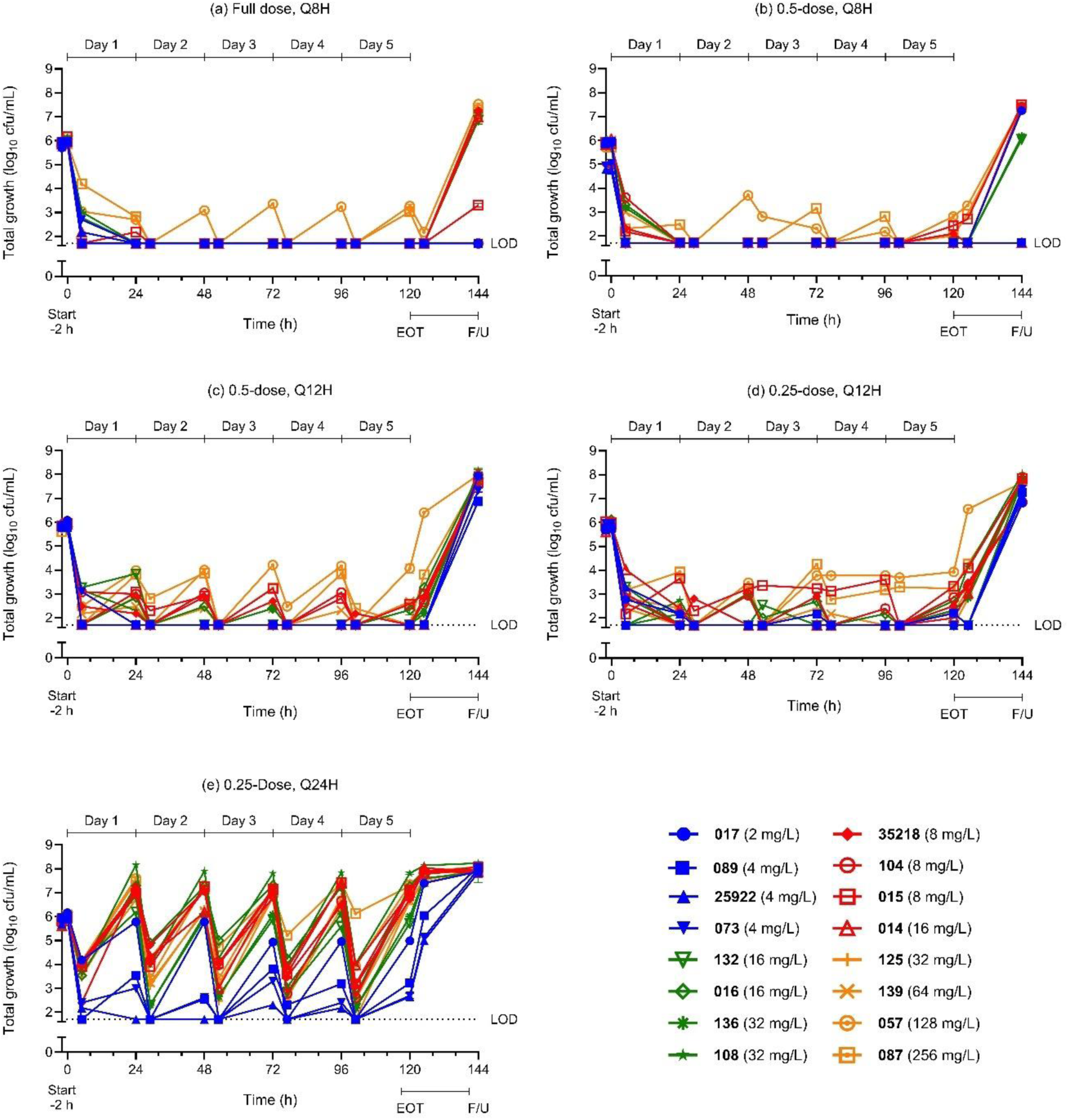
Growth response of *E. coli* uropathogens after exposure to simulated oral amoxicillin-clavulanic acid. Antibiotics were given for five days (120 h; end of treatment, EOT) and without antibiotics for a further 24 hours to detect any additional regrowth (144 h, Follow up, F/U). Five amoxicillin-clavulanic acid exposure experiments are shown. Limit of detection (LOD) was 50 cfu/ml (1.7 log_10_ cfu/ml).

**Table 2.** Change in bacterial density at end of treatment (EOT, 120 h) and follow up (F/U, 144 h).

| Isolate no. | Bacterial response after amoxicillin-clavulanic acid exposure (Δ log<sub>10</sub> cfu/mL) |  |  |  |  |  |  |  |  |  |
|---|---|---|---|---|---|---|---|---|---|---|
|  | End of treatment (EOT, 120 h) |  |  |  |  | Follow-up (F/U, 144 h) |  |  |  |  |
|  | Full-dose, q8 | 0.5x, q8 | 0.5x, q12 | 0.25x, q12 | 0.25x, q24 | Full-dose, q8 | 0.5x, q8 | 0.5x, q12 | 0.25x, q12 | 0.25x, q24 |
| ECO 017 | - | - | - | - | -1.2 | - | 1.4 | 1.9 | 0.9 | 1.7 |
| UTI 089 | - | - | - | -3.5 | -2.7 | - | - | 1.1 | 1.5 | 2.2 |
| ATCC 25922 | - | - | - | - | -3.3 | - | - | 1.1 | 1.2 | 1.9 |
| CFT 073 | - | - | - | - | -3.2 | - | - | 1.4 | 1.4 | 2.1 |
| ATCC 35218 | - | -2.8 | - | -3.3 | 0.7 | 1.3 | 2.5 | 1.8 | 1.0 | 1.8 |
| ECO 104 | - | - | - | -3.1 | 1.3 | - | 1.6 | 1.9 | 0.9 | 2.2 |
| ECO 015 | - | -3.4 | -3.4 | -2.7 | 1.2 | -2.9 | 1.7 | 1.7 | 1.8 | 2.1 |
| ECO 014 | - | - | - | -3.9 | 1.0 | 1.0 | - | 2.1 | 2.0 | 1.9 |
| ECO 132 | - | - | - | - | -0.3 | 0.9 | 0.3 | 2.0 | 1.8 | 1.8 |
| ECO 016 | - | - | -3.5 | -3.4 | 1.2 | - | 1.5 | 2.2 | 1.7 | 1.9 |
| ECO 136 | - | - | - | -3.6 | -0.1 | 1.3 | 0.4 | 1.9 | 1.8 | 1.7 |
| ECO 108 | - | - | - | -3.5 | 1.7 | 1.2 | - | 2.2 | 2.0 | 2.1 |
| ECO 125 | - | - | - | - | 0.8 | 1.1 | 1.5 | 1.9 | 2.1 | 2.1 |
| ECO 139 | - | - | -3.3 | -3.6 | 0.6 | - | 1.4 | 1.9 | 1.7 | 1.8 |
| ECO 057 | -2.61 | -3.1 | -1.9 | -2.1 | 1.3 | 1.7 | 1.5 | 2.1 | 1.6 | 2.0 |
| ECO 087 | -2.85 | -3.7 | - | -2.6 | 1.2 | 1.4 | 1.7 | 1.9 | 1.9 | 1.8 |

AMX, Amoxicillin. CLA, Clavulanic acid. CLA exposures were 1/10<sup>th</sup> of the amoxicillin exposure. A dash (-) indicates no growth detected. Limit of detection was 50 cfu/mL (1.7 log<sub>10</sub> cfu/ml). Target amoxicillin exposure for each experiment were as follow: Full-dose 8-hourly - AUC<sub>0-24</sub> 14272, C<sub>max</sub> 935; Half (0.5x) dose, 8-hourly (Q8H) - AUC<sub>0-24</sub> 7073, C<sub>max</sub> 467; Half-dose, 12-hourly (Q12H) - AUC<sub>0-24</sub> 4757, C<sub>max</sub> 440; Quarter (0.25x) dose, 12-hourly - AUC<sub>0-24</sub> 2379, C<sub>max</sub> 220; Quarter-dose, 24-hourly (Q24H) - AUC<sub>0-24</sub> 1189, C<sub>max</sub> 217.

### Amoxicillin only in the BIM

There were marked differences in the bacterial response following the half-dose, 8 hourly amoxicillin (without clavulanic acid) as compared to the same exposure with the addition of clavulanic acid. During the amoxicillin-only experiment, all isolates with a beta-lactamase gene displayed maximal growth by 24 hours, which was maintained for the total duration of therapy. Only isolates with no beta-lactamase gene (UTI089, ATCC 25922 and CFT073) had no detectable growth at EOT. In contrast, when the same amoxicillin exposure was combined with clavulanic acid, we observed reductions in bacterial density in all isolates during therapy, undetectable growth at EOT in 12/16 isolates, and the remaining 4 isolates had >2 log kill at EOT (Figure 5).

**Figure 5.**
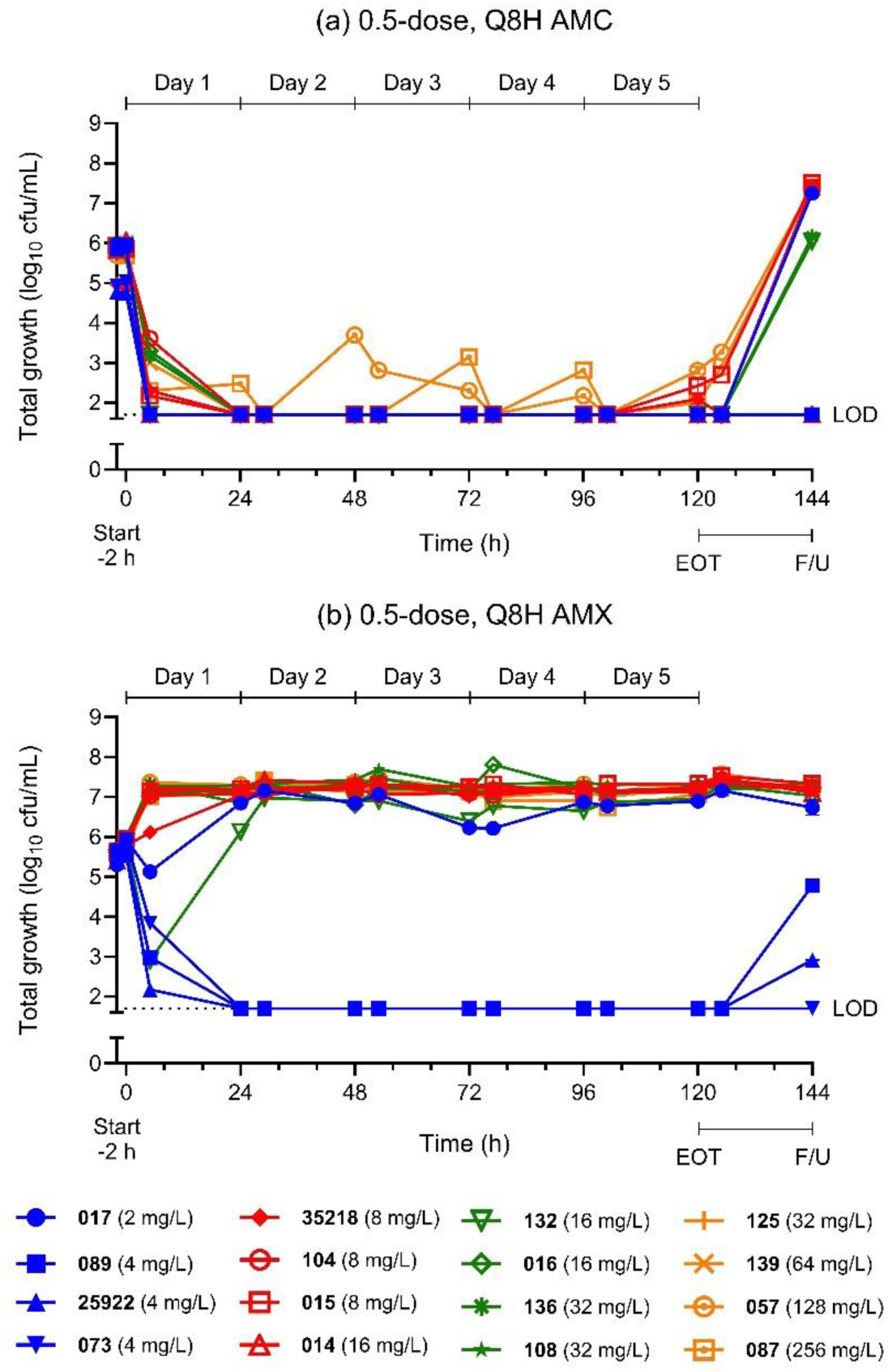
Amoxicillin-only versus amoxicillin-clavulanic acid exposure. Bacterial response in the bladder infection model following the half-dose amoxicillin-only 8-hourly experiment the compared with same exposure with the addition of clavulanic acid (1:10 ratio).

Beta-lactamase enzymatic hydrolysis of amoxicillin was demonstrated from the PK samples collected during the amoxicillin-only exposure experiment. Two sampled bladder compartments (ECO108 and ECO057) had undetectable amoxicillin concentrations at the day 5 expected peak concentration timepoint (Table S4). Supplementary analysis identified an additional amoxicillin degradation peak in the chromatogram at 1.6 min. The relative abundance of the degradation product, as a percentage of the chromatographic peak area, varied between strains. For ECO108 and ECO057 (with CTX-M-27 and TEM-1 genes) the degradation peak accounted for 100%. ECO015 (also with CTX-M-27 and TEM-1 genes) had a degradation peak of 17.4%. All other isolates had lower relative abundance of the degradation production. Isolates without an ESBL gene (UTI089, CFT073, ATCC 25922) had no degradation peak identified. In contrast with the corresponding amoxicillin-clavulanic acid experiment, the relative abundance of the amoxicillin degradation peak from samples collected at the same day 5 expected peak concentration time-point was the same for all bladder compartments (Figure S7 and S8).

### PK/PD analysis

The CLSI-MIC (2:1 amoxicillin-clavulanic acid ratio) based E_max_ model best describes the primary and secondary PD endpoints (Figure 6). For the change in bacterial density at EOT, the target %T_>MIC_ ranged from 52.7% for stasis up to 82.8% for 3 log_10_ kill (R^2^ 0.9062), and for AUC_0-24_/MIC, the target values ranged from 53.3 for stasis up to 285.8 for 3 log_10_ kill (R^2^ 0.8006) (Table 3). Similar targets were obtained for the secondary outcome that assessed the total bacterial response (AUBKC_0-144_) with improved goodness-of-fit R^2^ values for %T_>MIC_ and AUC_0-24_/MIC (0.9391 and 0.8904, respectively). EUCAST-MIC (fixed concentration of 2 mg/L clavulanic acid) displayed a poorer goodness-of-fit (R^2^ 0.80-0.82) based analyses (Figure S9 and Table S5). A greater goodness-of-fit of the E_max_ curve was observed using MICs measured in mSHU for the %T_>MIC_ analysis (R^2^ 0.96-0.97; Figure S10, Table S6).

**Figure 6.**
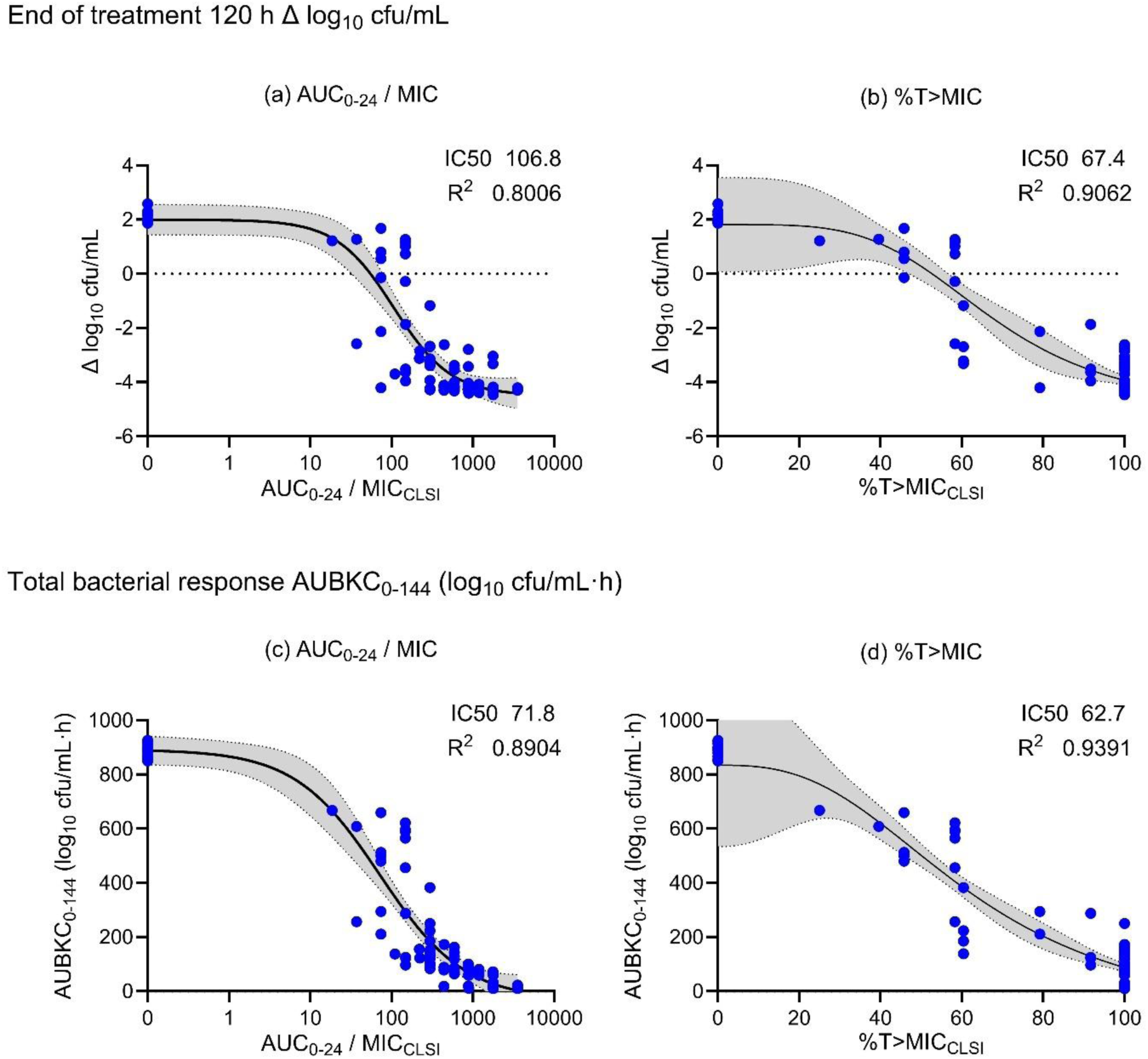
Exposure-response relationships. Applying baseline amoxicillin-clavulanic acid MIC values using CLSI methodology (2:1 amoxicillin-clavulanic acid ratio). The four-way parametric E_max_ models above detail the primary (end of treatment Δ log_10_ CFU/mL) and secondary outcomes (AUBKC_0–144_). Graphs have not been constrained. The shaded area indicates the 95% confidence band. The 50% maximal inhibitory concentration (IC_50_) and coefficient of determination (R^2^) are reported.

**Table 3.** Amoxicillin-clavulanic acid urinary PK/PD targets.

|  | **%T>MIC<sub>CLSI</sub> (95% CI)** | **AUC<sub>0-24</sub> /MIC<sub>CLSI</sub> (95% CI)** |
|---|---|---|
| **End of treatment outcome (at 120 h; Δ log<sub>10</sub> cfu/mL)** |  |  |
| Stasis | 52.7% (47.1 - 56.7%) | 53.3 (32.4 - 78.0) |
| 1 log<sub>10</sub> kill | 61.8% (58.0 - 67.1%) | 94.4 (65.9 - 119.8) |
| 2 log<sub>10</sub> kill | 71.1% (65.8 - 78.9%) | 156.9 (123.3 - 197.0) |
| 3 log<sub>10</sub> kill | 82.8% (74.2 - 89.7%) | 285.8 (218.7 - 381.4) |
| R<sup>2</sup> | 0.9062 | 0.8006 |
| **Total bacterial response (AUBKC<sub>0-144</sub>; log<sub>10</sub> cfu/mL·h)** |  |  |
| Stasis | 48.8% (43.2 - 52.6%) | 45.8 (27.3 - 62.9) |
| 1 log<sub>10</sub> reduction | 59.2% (55.5 - 63.7%) | 85.8 (62.7 - 107.1) |
| 2 log<sub>10</sub> reduction | 71.1% (65.6 - 78.2%) | 165.3 (133.2 - 204.5) |
| 3 log<sub>10</sub> reduction | 87.1% (78.5 - 92.4%) | 370.5 (286.4 - 481.7) |
| R<sup>2</sup> | 0.9391 | 0.8904 |

### Monte Carlo simulations

Applying variability in expected human urinary concentration (± 50%) and assuming urinary exposure ratio of amoxicillin to clavulanic acid is maintained at least at a 10:1 ratio, MCS indicated that 500 mg amoxicillin administered every 8-hours would achieve all EOT %T_>MIC_ targets (up to a 3 log_10_ kill) in >95% of virtual patients infected with *E. coli* isolates with MICs ≤ 64 mg/L, regardless of MIC method (CLSI or EUCAST). With less frequent dosing, 500 mg and 250 mg 12-hourly, to achieve all EOT %T_>MIC_ targets in >95% of virtual patients require isolates with CLSI MICs of ≤ 16 mg/L and ≤ 8 mg/L, and EUCAST MICs of ≤ 32 mg/L and ≤ 16 mg/L, respectively. If an EOT *in vitro* PD target of 1 log_10_ kill is applied, then >95% of virtual patients can be achieved for isolates with MIC ≤ 128 mg/L for 500 mg 8-hourly, MIC ≤ 64 mg/L for 500 mg 12-hourly, and MIC ≤ 32 mg/L for 250 mg 12-hourly, regardless of MIC method (Figure 7 and S11).

**Figure 7.**
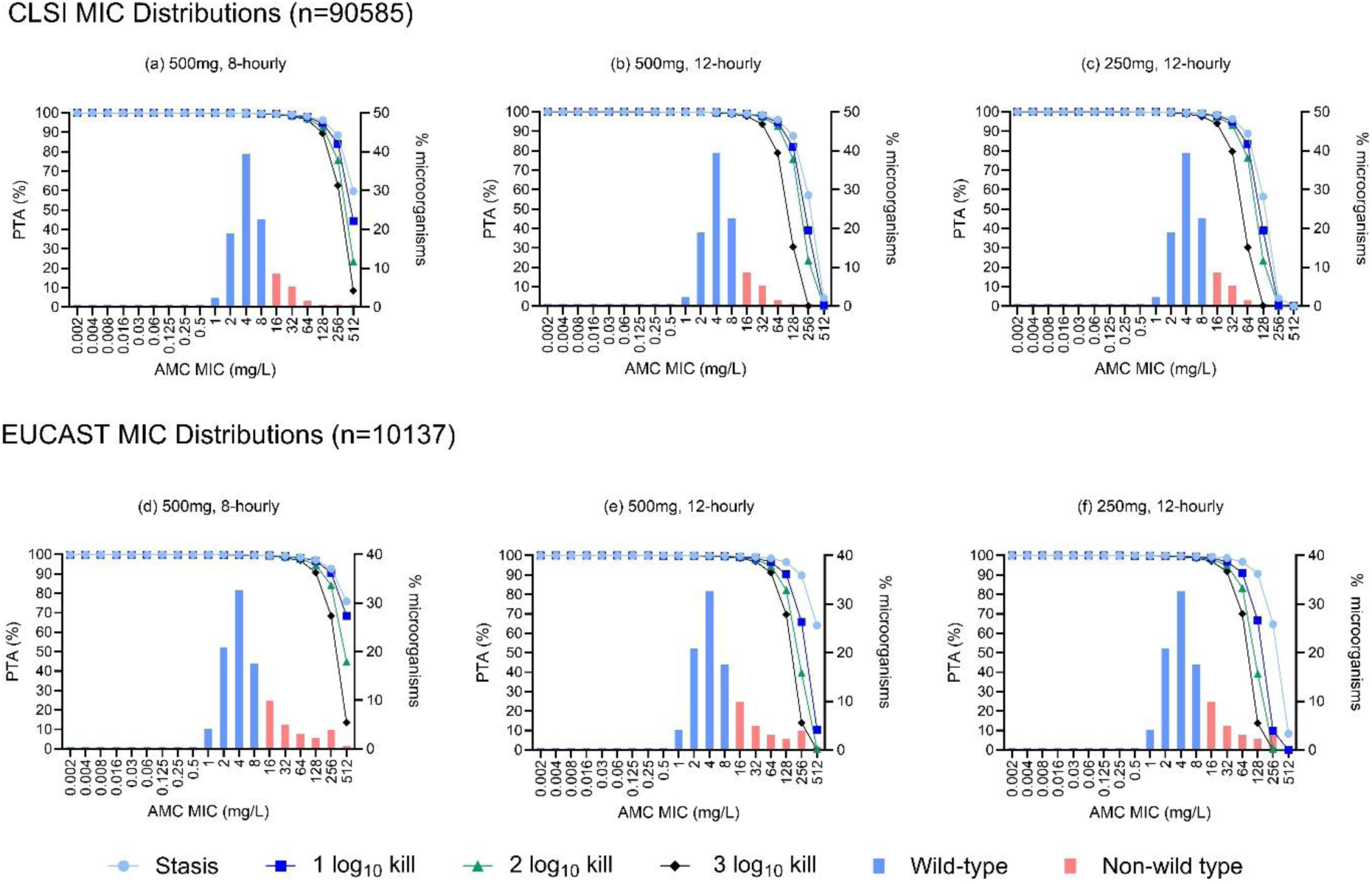
Monte Carlo simulations. Simulation of expected urinary exposure following (a,d) 500 mg amoxicillin acid given 8 hourly or (b,e) 12 hourly, and (c,f) 250 mg amoxicillin given 12 hourly for 5,000 virtual patients. Clavulanic acid exposure assumed to be one-tenth of the amoxicillin exposure. Graphs show probability of target attainment for %T_>MIC_ targets at end-of-treatment (120 h) corresponding to bacterial stasis and log killing, as determined in this study. Graphs overlay CLSI *E. coli* amoxicillin-clavulanic acid MIC distributions (top row; provided by J. Turnidge [unpublished]) and EUCAST MIC distributions (bottom row; from the EUCAST

## DISCUSSION

Using a dynamic bladder infection *in vitro* model simulating human urinary pharmacokinetics, we demonstrate rapid and sustained killing of ESBL-producing *E. coli* by amoxicillin-clavulanic acid. Killing at end of treatment was observed in both wild type and non-wild type isolates and was maintained when simulated exposure and dosing frequency were reduced. Killing was time-dependent, and %T>_MIC_ was the PK/PD index most closely associated with efficacy; approximately 50%, 60%, 70% and 80% %T>_MIC_ required for stasis, 1, 2, and 3 log_10_ kill, respectively. Monte Carlo simulation indicated that 500/125 mg three times daily achieves higher target attainment compared to twice-daily dosing, with a high likelihood treatment success against isolates with an MIC up to 64 mg/L, irrespective of whether MICs were determined by CLSI or EUCAST methodology. These data establish the urinary exposure-response relationship for amoxicillin-clavulanic acid against ESBL-producing *E. coli* and inform the dosing and susceptibility conditions under which treatment could be expected to succeed.

In humans, few studies have investigated the utility of amoxicillin-clavulanic acid in treating UTIs caused by ESBL-producing uropathogens. In a small retrospective clinical study, 15 patients with cystitis caused by ESBL-producing *E. coli* received amoxicillin-clavulanic acid oral therapy after susceptibility was confirmed (39). All 15 cystitis patients displayed clinical cure. All isolates in this study, however, had an amoxicillin-clavulanic acid MIC of ≤ 8 mg/L, so it is uncertain whether these findings can be extrapolated to the higher breakpoints for uncomplicated UTIs outlined by CLSI and EUCAST. The same study additionally showed the utility of oral amoxicillin-clavulanic acid in complicated UTIs as well, with 8/9 patients with ESBL-positive *E. coli* UTIs causing vesicoureteral reflux-cystitis and hyperactive bladder-cystitis reaching clinical cure (39). In another retrospective cohort, amoxicillin-clavulanic acid was found to be non-inferior to standard of care in treating ESBL-producing uropathogens in pyelonephritis cases, with amoxicillin-clavulanic acid displaying a 19.2% failure rate as compared to the 30.3% failure rate with standard of care, although no MICs were reported (40). The largest study of amoxicillin-clavulanic acids utility in the treatment of uncomplicated UTIs to date, comes from the REASSURE trial (41). In this study, sulopenem was compared to the oral amoxicillin-clavulanic acid combination (875/125mg, bidaily), and sulopenem was found non-inferior to amoxicillin-clavulanic acid in treating uncomplicated UTIs (76.1% clinical cure vs 76.5% clinical cure). Interestingly, this study also observed that in ceftriaxone and amoxicillin-clavulanic acid-resistant isolates, out of 70 infection, 56 (80%) still showed clinical response to the combination. These studies, in conjunction with the dependable bacterial kill observed in our *in vitro* model, support the use of amoxicillin-clavulanic acid for treating UTIs caused by ESBL-producing, ceftriaxone-resistant *E. coli* uropathogens.

We observed the best goodness-of-fit in the exposure-response analysis when using amoxicillin-clavulanic acid MICs measured by CLSI-methodology compared with EUCAST. CLSI applies a 2:1 amoxicillin to clavulanic acid ratio for MIC determination, whilst EUCAST recommends a fixed concentration of clavulanic acid (2 mg/L). EUCAST specifically provides oral breakpoints for uncomplicated UTIs (S ≤ 32 mg/L, R > 32 mg/L). Whereas CLSI breakpoints are reported as: S ≤ 8/4 µg/mL, I^ = 16/8 µg/mL, R ≥ 32/16 µg/mL), including the comment that when oral amoxicillin-clavulanic acid is used the breakpoint is only for therapy of uncomplicated UTIs or for completion of therapy for systemic infection. Furthermore, the “intermediate” category indicates that amoxicillin-clavulanic acid has the potential to concentrate in the urine, which may or may not be incorporated into clinical laboratory reporting. This has contributed to variable agreement between the two MIC methodologies, especially with ESBL-producing organisms (42, 43). Our data has shown that the higher concentrations of clavulanic acid used with the CLSI methodology results in lower MICs in beta-lactamase-producing organisms. The EUCAST methodology tends to yield greater range and higher MICs.

There is wide variability in both amoxicillin and clavulanic acid excretion and observed urinary concentrations after oral dosing (44, 45). In a 1972 paper (46), the average concentration of amoxicillin in pooled urine collected over the six-hour period after dosing was 580 µg/mL (range 50 - 1,600 µg/mL) for the 250 mg dose and 1,100 µg/mL (range 115 - 1,850 µg/mL) for the 500 mg dose. Neither urine volume nor the number of subjects was reported, precluding calculation of a coefficient of variation. Notably, the authors separately report a comparatively narrow range for the proportion of the dose excreted in urine over six hours (58 - 68%), suggesting that variable urine output, rather than variable amoxicillin absorption and renal excretion, contributes to the observed spread in measured concentrations. The full-dose exposure experiment we ran in the BIM generated a clinically plausible urinary exposure profile following a 500 mg oral dose of amoxicillin, correcting for the fact that 100% of the input antibiotic dose is delivered to the bladder compartment. Clavulanic acid urinary exposure is less certain due to few studies investigating urinary exposure. Urinary recovery for clavulanic acid is high albeit also highly variable, with one study finding an average urinary concentration after an oral dose of 125 mg of clavulanic acid (co-formulated with amoxicillin) being 426 mg/L (53-926 mg/L) over the first four hours, and 43.4 ± 15.6% of the oral dose recovered in urine after 24 hours (47). The clavulanic acid exposure we have targeted are lower than that reported in the literature. Only a small amount of clavulanic acid is required to inhibit beta-lactamases (<1 mg/L) (48, 49), so even assuming concentrations at the lower end of that spectrum, we would expect sufficient inhibition in a human patient.

We demonstrated that high urinary amoxicillin concentrations alone were insufficient to kill ESBL-producing strains. The amoxicillin-only exposure experiment, matched to the amoxicillin-clavulanic acid dosing exposure, confirmed that bacterial kill of ESBL-producing isolates was driven by clavulanic acid restoring the activity of amoxicillin, and not due to the high concentrations of amoxicillin. Degradation products of amoxicillin were detected when amoxicillin was administered without the protection of clavulanic acid, demonstrating critical inhibition of ESBL enzymatic activity by clavulanic acid in the urinary environment.

Following antibiotic exposure in the model, we observed regrowth without emergence of resistance. This has been similarly reported following the testing of other antibiotics in the BIM (33, 50, 51). A possibility is that following antibiotic exposure in a nutritionally-deplete environment, such as the mSHU media, the bacteria are evolutionally primed to become metabolically inactive leading to bacterial persistence (52). Such persistent subpopulations of bacteria in a low-metabolic or a near-dormant state can then survive bactericidal antibiotic concentrations, whilst remaining phenotypically susceptible (53, 54). Infections isolated from UTI patients can display persistent subpopulations (55), but their clinical relevance remains poorly defined. As no immune cells are present in the BIM to potentially clear these persister cells, it is uncertain if the bacterial regrowth we observed after antibiotic exposure is of clinical importance, or an artefact of our *in vitro* model.

There have been recent therapeutic advancements for the treatment of UTIs, with three new agents having recently received FDA approval: sulopenem etzadroxil-probenecid (41, 56), gepotidacin (57, 58), and tebipenem pivoxil (59–61). The FDA also recently approved pivmecillinam (200 mg three times per day)(62), which is a lower dose compared with the IDSA

Guidance recommendation for the higher 400 mg three times per day dosing (29). Considering existing oral agents, in a 2026 randomized clinical trial, the SCOUT study confirmed nitrofurantoin (100 mg three times per day for 5-days) was the most effective first-line treatment option for women with uncomplicated UTIs, compared with fosfomycin (single 3g dose; two 3g doses given daily), and pivmecillinam (400 mg three times per day 3-days) (63). Amoxicillin-clavulanic acid was not included in this study.

The strength of the BIM in PK/PD assessments is the generation of human urinary antibiotic exposures together with a simulation of urodynamics within a synthetic and reproducible urinary liquid environment. The model, however, lacks an immune system and other physiological features that significantly factor in therapeutic success. This includes testing at atmospheric conditions and absence of uroepithelial cells of the bladder wall, which provides a structure for bacteria to adhere to and infiltrate (64, 65). Additionally, each independent run of the bladder infection model was only conducted once without replication, so some biological variability may affect results.

Human urine has intrinsically complex chemistry that is difficult to replicate with a synthetic alternative. We have shown that mSHU provides a standardized and robust substitute for urine (66), but it is not a perfect mimic. We observed that AMC MICs were lower in mSHU compared to pooled human urine, which may be due to chemical and pH differences, yet drug-free growth kinetics were matched. Despite this, the translatability of our findings is maintained, considering that CLSI-derived MICs were less than a doubling dilution lower in mSHU compared with urine.

The PK/PD index for clavulanic acid was not independently characterized, and the MCS assumed clavulanic acid urinary exposure was maintained at a ratio of at least one-tenth of the amoxicillin exposure. This was due to the use of a parenteral formulation of amoxicillin-clavulanic acid with a 10:1 ratio. Whilst this ratio is physiologically plausible, we did not specifically test different ratios of clavulanic acid, and our analyses assume that clavulanic acid concentrations remain sufficient to restore the activity of amoxicillin in all experiments. The literature suggests that activity of clavulanic acid is best described by time above a concentration threshold (67, 68). As evidenced by the amoxicillin-only experiment, any bacterial kill against ESBL-producing isolates in the BIM would indicate that there was sufficient concentration of clavulanic acid to restore the activity of amoxicillin. Therefore, it is less likely that regrowth observed at low simulated exposures could be a consequence of the clavulanic acid threshold not being met.

Using a dynamic bladder infection model, we present *in vitro* PK/PD data that argues against a blanket exclusion of amoxicillin-clavulanic acid for uncomplicated UTI caused by ESBL-producing *E. coli*, as currently suggested in the IDSA 2026 Guidance (29). Accounting for the expected variability in urinary amoxicillin concentrations after oral administration, and assuming urinary clavulanic acid concentrations of at least one-tenth those of amoxicillin, our exposure-response profiling supports uncomplicated UTI-specific breakpoints for a 5-day course of amoxicillin-clavulanic acid: MIC ≤64 mg/L (CLSI or EUCAST) with 500/125 mg three times daily, and MIC ≤16 mg/L (CLSI) or ≤32 mg/L (EUCAST) with 500/125 mg twice daily. Our data establish a pharmacological rationale for revisiting amoxicillin-clavulanic acid in this indication under appropriately defined susceptibility criteria.

## ACKNOWLEDGEMENTS

This study was funded by the US Food and Drug Administration (FDA) under contract number 75F40123C00185. Views expressed are solely the responsibility of the authors and do not necessarily represent the official views of the FDA. C.R.B. Anderson was supported by an Australian Government Research Training Program (RTP) Scholarship. I.J. Abbott has undertaken advisory board consultancy for Merck Sharp & Dohme (Australia) (June 2021), GSK (November 2023), and Clinigen/Shionogi (May 2026), and invited speaker for Coloplast (August 2026) and Becton Dickinson (February 2025, February 2026), no personal remuneration received. I.J. Abbott received support from an Australian National Health and Medical Research Council (NHMRC) Emerging Leader-1 Investigator Grant. A.Y. Peleg has undertaken advisory board consultancy for Clinigen/Shionogi. J.A. Roberts would like to acknowledge funding from the Australian National Health and Medical Research Council for a Centre of Research Excellence (APP2007007) and an Investigator Grant (APP2009736). All other authors: none to declare.

We thank Dr Adam Jenney, Microbiology Unit, Department of Pathology, Alfred Health, Melbourne, Australia for providing the baseline collection of *E. coli* clinical isolates. We thank Jenny Ordonez for assistance with antibiotic quantification. We thank Professor John Turnidge for providing the *E. coli* amoxicillin-clavulanic acid MIC distribution data with both EUCAST and CLSI methodology. We would additionally like to thank the members of the US FDA who contributed to the review of the manuscript: Thushi Amini, Rebecca Dean, Jalal Sheikh, Elizabeth-Story Roller, Zhixia (Grace) Yan Danielsen, Avery Goodwin, Anthony Nicasio, Peter Kim, Dmitri Iarikov, and Ramya Gopinath.

## SUPPLEMENTARY MATERIAL SUPPLEMENTARY METHODS

### Antibiotic stock solutions & antibiotic-impregnated agar

Anhydrous amoxicillin powder (Merck Germany, A8523) was dissolved in dimethyl sulfoxide (DMSO; Merck Germany, 472301) solvent and deionized water as a diluent to develop an amoxicillin stock solution. Final DMSO concentrations were ≤ 0.5% when performing broth microdilution. For dissolving potassium clavulanate (Merck Germany, 3345) and lithium clavulanate (Merck Germany, L0720000) powder, phosphate buffer (pH 6.0, 0.1 mol/L) was utilized as both solvent and diluent. When preparing MHA supplemented with amoxicillin-clavulanic acid (AMC-CAMHA-32/2 mg/L), agar was allowed to cool to 45°C before the addition of amoxicillin and clavulanic acid, then stirred. Plates were poured and allowed to cool to room temperature. Plates were stored at 4-8°C until use.

### Modified synthetic human urine (mSHU)

The makeup of modified synthetic human urine (mSHU), as well as the concentrations of each component, is listed below in Table S1. The original recipe for SHU (1) was modified by the addition of two growth supplements (0.2 g/L Bacto™ casamino acids [Thermofischer Scientific, USA] and 0.1 g/L yeast extract [Condalab, Spain]). Ultrapure water was heated, and all chemicals were solubilized stepwise in the order outlined in Table S1. Hydrochloric acid and sodium hydroxide were utilized to adjust the final pH to 5.6. Media was sterilized by pressure filtration (Millipore® Steripak Pressure Filter; Merck, Germany).

### Pooled Human Urine

Human urine was collected under ethics approval (project ID #27033). Criteria for urine donation excluded those who had taken antibiotics within the last month, displayed symptoms of a UTI, were pregnant or had a history of kidney disease. Upon collection, urine was pooled together by equal volumes from each volunteer, and filtered through a 0.2 µm polyethersulfone (PES) filter (Rapid Flow [Thermofischer Scientific, USA]). The sterile, pooled human urine was kept refrigerated at 2 - 8°C and utilized within 48 hours of collection. Two urine collections were performed, the first batch was collected from twelve healthy volunteers (nine female, three male), which was utilized for the growth kinetics assay and amoxicillin alone BMD MIC. The resulting pooled human urine displayed a pH of 6.23. The second batch of urine was collected from twelve healthy volunteers (eleven females, one male), which was utilized for the amoxicillin-clavulanic acid BMD MIC (both EUCAST and CLSI-derived methodology). The resulting pooled human urine displayed a pH of 6.91.

### Bladder infection model

The liquid medium (mSHU) was run at a flow rate of 400 mL/h from fresh media reservoirs through the intestinal compartment to the circulatory compartment. For the duration of each experiment, the volume of the intestinal compartment was maintained at 350 mL, and the volume of the circulatory compartment was maintained at 1000 mL. A multichannel pump then delivered media from the circulatory compartment to the sixteen bladder compartments run in parallel. The inflow rate to each bladder was 25 mL/h, with bladder compartments heated within a water bath held at 37°C and internal mixing via magnet stirrers. Media with each bladder compartment was allowed to increase in volume before voiding via a timed void schedule pump every four hours. The post-void residual was 7.1 ± 0.9 mL. Bladders were inoculated with 10 mL of 10^6^ CFU/mL at T-2, to correlate with the total bacteria count expected in human UTIs (≥10^5^ CFU/mL in a 200 mL void). Inoculated bacteria were sourced from overnight cation-adjusted Mueller-Hinton agar (CAMHA) growth cultures, subculture from frozen stock. No pre-treatment to induce or select for ESBL production was performed. Automated dosing of antibiotics was controlled through two pumps, one which empties the content of the intestinal compartment and one that supplies a new dose of antibiotics. Their activity was controlled via wired connection to a timed switch control unit, which was programmed to dose antibiotics automatically every 8, 12 or 24 hours.

### Quantification of bacterial density

Samples were collected directly from each bladder compartment to assess changes in bacterial density during each experiment in the BIM. Each 1 mL sample was pre-processed to prevent antibiotic carryover. Samples were centrifuged at 10,000 g for 5 minutes to pellet the bacteria present. 900 µL of supernatant was removed and fresh 900 µL of 0.85% saline was added to wash the sample of antibiotics. This process was repeated twice. Samples then underwent a 10-fold serial dilution, with 20 μL of each dilution being plated on agar (CAMHA or AMC-CAMHA-32/2 mg/L) and incubated aerobically at for 35 ±1°C for 18-24 h before colony counts were performed. The AMC-CAMHA-32/2 mg/L were incubated for an additional 24 hours after an initial count to detect any slower growing colonies. The final concentrations of antibiotics in the supplemented agar (32 mg/L of amoxicillin and 2 mg/L of clavulanic acid) were chosen to correlate with the EUCAST uncomplicated UTI breakpoints for amoxicillin-clavulanic acid oral therapy. Growth on this plate was used as a screening method to identify emergence of resistance or quantification of a resistant sub-population, and not as a measurement of MIC. The limit of detection (LOD) was 50 CFU/mL (1.7 log_10_ CFU/mL). Fine haze, pinpoint or single colonies were not considered true growth and not counted.

### Antibiotic concentration quantification

Antibiotics samples were taken from 3 representative bladder compartments across set timepoints from 24 h to 104 h to ascertain antibiotic levels. Sampling times correlated to C_min_ and C_max_. On the fifth day of sampling, additional samples were taken to capture concentrations before and after the C_max._ Additionally, the C_max_ sample on the fifth day was taken from all 16 bladder compartments to capture inter-compartment variability. Samples taken were filtered (PES membrane, 0.22μm) and stored at -80°C until antibiotic levels were quantified by ultra-high-performance liquid chromatography-tandem mass spectrometry (UHPLC-MS/MS). Supplementary HPLC-PDA was performed to analyze peak areas for amoxicillin and amoxicillin-degradation products, with the degradation peak relative abundance reported as a percentage of the chromatographic peak area. Stability of amoxicillin and clavulanic acid in mSHU at different temperatures was also assessed. Amoxicillin and clavulanic acid were suspended in mSHU (1000:100mg/L), and stored at three temperatures; 4-8°C, 20-22°C and 37°C. This was done to mimic the conditions of the drug reservoir, circulatory compartment, and bladder compartment respectively. Samples were taken at baseline, 8 hours, and 24 hours.

### Whole genome sequencing

Genomic DNA was extracted using Genfind v3 kit (Beckman Coulter). Libraries were prepared using the Nextera Flex DNA Library Prep Kit (Illumina) as per manufacturer’s instruction, and 150 bp paired-end sequencing was performed on the NovaSeq 6000 system (Illumina). Illumina reads were assembled using unicycler v0.5.0 (2) with default parameters. One isolate (ECO 108) that was sequenced on ONT platforms after failing. ONT sequencing was assembled using a custom assembly pipeline (3). Briefly, the pipeline filters read using Filtlong v0.2.1 (4) with the parameters --min_length 1000 and --keep_percent 95, assembles using Flye v2.9 (5) with the --nano-hq tag, and polishes using Medaka v1.7 (6). MLST was performed using mlst v2.23.0 (7) with the *E. coli* scheme. AMR gene detection was performed using AMRFinderPlus v4.0.15 (8) with *E. coli* set as the organism.

## SUPPLEMENTARY TABLES

**Table S1.**
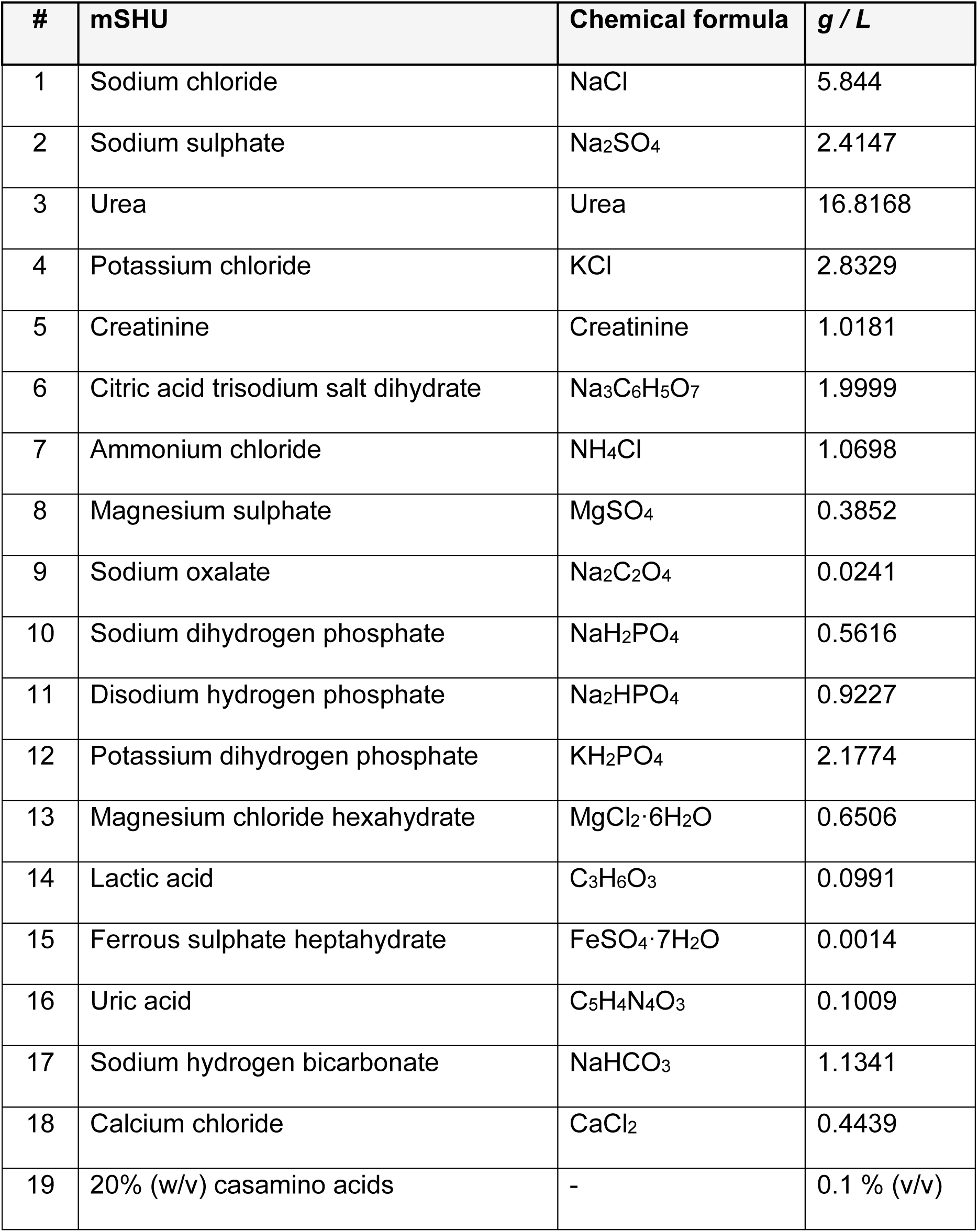
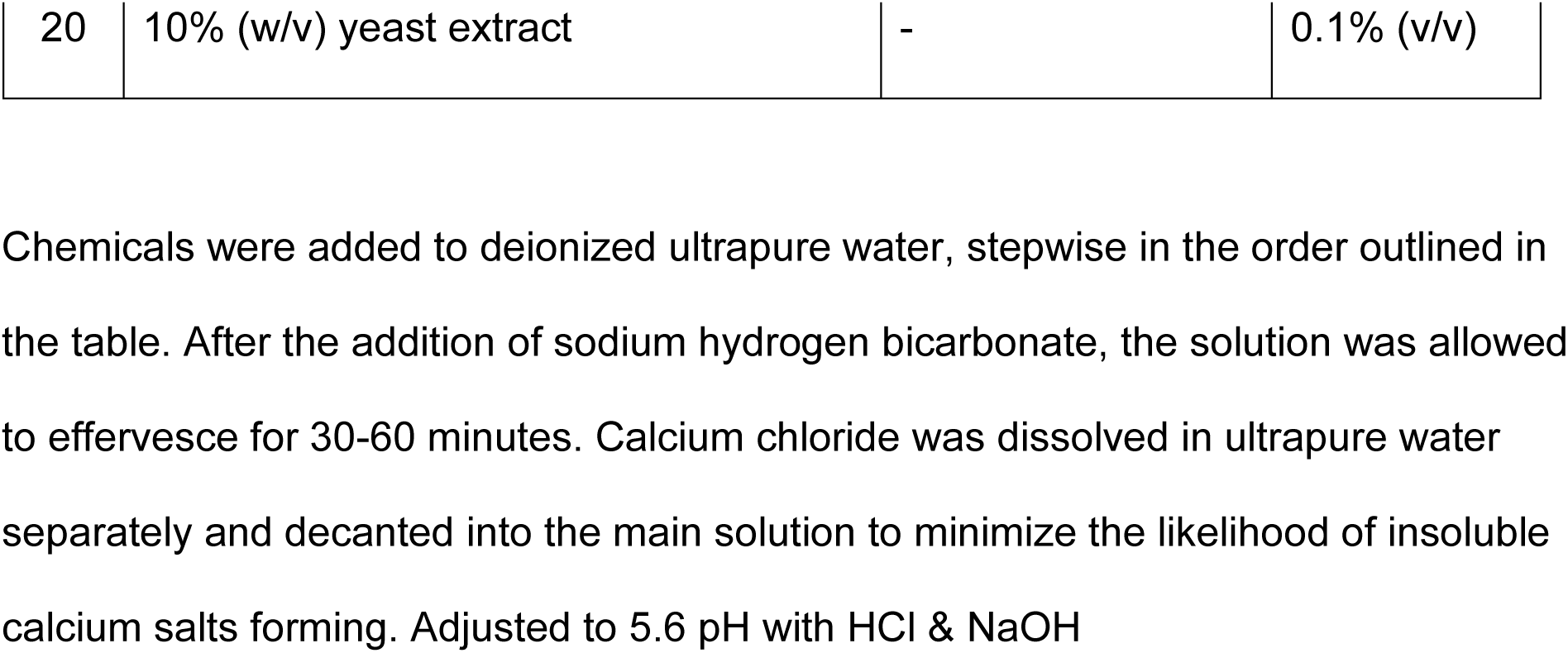
Modified synthetic human urine formulation.

**Table S2.** Amoxicillin MICs of sixteen *E. coli* tested in the bladder infection model.

| **Isolate no.** | **ST** | **Beta-lactamase gene** | **Baseline AMC BMD MIC (median (range), mg/L)** |  |  |
|---|---|---|---|---|---|
|  |  |  | **CAMHB<sup>a</sup>** | **mSHU<sup>a</sup>** | **Human urine<sup>b</sup>** |
| **ECO 017** | 219 | CTX-M-15 | >256 (>256) | 128 (128 - >256) | >256 |
| **UTI 089** | 95 | - | 4 (4) | 2 (1 - 2) | 2 |
| **ATCC 25922** | 73 | - | 4 (4) | 1 (0.5 - 2) | 4 |
| **CFT 073** | 73 | - | 4 (4 - 8) | 1 (1 - 2) | 2 |
| **ATCC 35218** | 127 | TEM-1 | >256 (>256) | >256 (>256) | >256 |
| **ECO 104** | 95 | CTX-M-55, TEM-1 | >256 (>256) | >256 (>256) | >256 |
| **ECO 015** | 131 | CTX-M-27, TEM-1 | >256 (>256) | >256 (>256) | >256 |
| **ECO 014** | 38 | CTX-M-14 | >256 (>256) | >256 (>256) | >256 |
| **ECO 132** | 648 | CTX-M-65, OXA-10 | >256 (>256) | 128 (64 - 128) | >256 |
| **ECO 016** | 131 | CTX-M-14, TEM-1 | >256 (>256) | >256 (>256) | >256 |
| **ECO 136** | 131 | CTX-M-15, OXA-1 | >256 (>256) | >256 (>256) | >256 |
| **ECO 108** | 393 | CTX-M-27, TEM-1 | >256 (>256) | >256 (>256) | >256 |
| **ECO 125** | 131 | CTX-M-15, OXA-1 | >256 (>256) | >256 (>256) | >256 |
| **ECO 139** | 131 | CTX-M-15, OXA-1 | >256 (>256) | >256 (>256) | >256 |
| **ECO 057** | 131 | CTX-M-27, TEM-1 | >256 (>256) | >256 (>256) | >256 |
| **ECO 087** | 1193 | CMY-2, CTX-M-15 | >256 (>256) | >256 (>256) | >256 |

<sup>*a*</sup> Triplicate BMD testing, MIC is given as median (range). <sup>*b*</sup> Single test performed.

**Table S3.** Antimicrobial disk diffusion testing for ampicillin and amoxicillin-clavulanic acid.

| **Isolate** | **AMP, 10 ug (mm)** | **EUCAST Interp.** | **CLSI Interp.** | **AMC, 30 ug (mm)** | **EUCAST Interp.** | **CLSI Interp.** |
|---|---|---|---|---|---|---|
| **ECO 017** | 6 | R | R | 22 | S | S |
| **UTI 089** | 20.5 | S | S | 22 | S | S |
| **ATCC 25922** | 19.5 | S | S | 21 | S | S |
| **CFT 073** | 20.5 | S | S | 22 | S | S |
| **ATCC 35218** | 6 | R | R | 21 | S | S |
| **ECO 104** | 6 | R | R | 21.5 | S | S |
| **ECO 015** | 6 | R | R | 22 | S | S |
| **ECO 014** | 6 | R | R | 21 | S | S |
| **ECO 132** | 6 | R | R | 22 | S | S |
| **ECO 016** | 6 | R | R | 21 | S | S |
| **ECO 136** | 6 | R | R | 17 | S | I |
| **ECO 108** | 6 | R | R | 19 | S | S |
| **ECO 125** | 6 | R | R | 17 | S | I |
| **ECO 139** | 6 | R | R | 15.5 | S | I |
| **ECO 057** | 6 | R | R | 13.5 | R | I |
| **ECO 087** | 6 | R | R | 11 | R | R |

AMP = Ampicillin, AMC = Amoxicillin-clavulanic acid. Zone measurements are reported in millimeters. Zone measurements above are an average of two duplicate disc diffusion tests. EUCAST interpretations are AMP – S ≥ 14mm, R < 14mm; AMC (oral, uncomplicated UTI only) – S ≥ 16mm, R < 16mm. CLSI interpretations are AMP – S ≥ 17mm, I = 14-16mm, R ≤13 mm; AMC – S ≥18mm, I = 14-17mm, R ≤13 mm. A measurement of 6 mm indicates no measurable zone.

**Table S4.** Amoxicillin concentrations at day 5 peak concentration during the amoxicillin only experiment.

| **Isolate** | **AMX MIC** | **Beta-lactamase** | **AMX conc. (mg/L)** | **% AMX degradation peak** |
|---|---|---|---|---|
| **ECO 017** | >256 (>256) | CTX-M-15 | 400.9 | 6.9% |
| **UTI89** | 4 (4) | - | 460.6 | 0% |
| **ATCC 25922** | 4 (4) | - | 413.5 | 0% |
| **CFT073** | 4 (4 - 8) | - | 433.3 | 0% |
| **ATCC 35218** | >256 (>256) | TEM-1 | 358.0 | 13.1% |
| **ECO 104** | >256 (>256) | CTX-M-55, TEM-1 | 352.8 | 14.8% |
| **ECO 015** | >256 (>256) | CTX-M-27, TEM-1 | 320.3 | 17.4% |
| **ECO 014** | >256 (>256) | CTX-M-14 | 387.5 | 8.0% |
| **ECO 132** | >256 (>256) | CTX-M-65, OXA-10 | 357.7 | 11.2% |
| **ECO 016** | >256 (>256) | CTX-M-14, TEM-1 | 368.2 | 12.4% |
| **ECO 136** | >256 (>256) | CTX-M-15, OXA-1 | 405.3 | 3.1% |
| **ECO 108** | >256 (>256) | CTX-M-27, TEM-1 | 0.0 | 100% |
| **ECO 125** | >256 (>256) | CTX-M-15, OXA-1 | 378.8 | 3.2% |
| **ECO 139** | >256 (>256) | CTX-M-15, OXA-1 | 396.5 | 7.6% |
| **ECO 057** | >256 (>256) | CTX-M-27, TEM-1 | 0.0 | 100% |
| **ECO 087** | >256 (>256) | CMY-2, CTX-M-15 | 429.4 | 4.0% |

All 16 bladder compartments were sampled at day 5 expected peak concentration time-point and the amoxicillin concentrations quantified. The expected concentration was 467.4 mg/L. The HPLC-PDA peak analysis determined the relative contribution of the amoxicillin degradation products as a percentage, described in the table as the % AMX degradation peak. Note that there were additional small peaks not classified as a degradation peak, therefore the area associated with amoxicillin and degradation peak percentages do not always sum to 100% (Figure S6).

**Table S5.** Amoxicillin-clavulanic acid urinary PK/PD targets with EUCAST MICs.

|  | **%T>MIC<sub>EUCAST</sub> (95% CI)** | **AUC<sub>0-24</sub> /MIC<sub>EUCAST</sub> (95% CI)** |
|---|---|---|
| **End of treatment outcome (at 120 h; Δ log<sub>10</sub> cfu/mL)** |  |  |
| Stasis | 32.8% (~ - 43.5%) | 6.5 (1.6 - 15.8) |
| 1 log<sub>10</sub> kill | 47.6% (36.9 - 55.7%) | 18.9 (5.9 - 34.0) |
| 2 log<sub>10</sub> kill | 61.6% (53.2 - 72.2%) | 53.3 (27.5 - 87.5) |
| 3 log<sub>10</sub> kill | 77.9 (66.9% - 87.6%) | 176.6 (100.5 - 319.1) |
| R<sup>2</sup> | 0.7241 | 0.7493 |
| **Total bacterial response (AUBKC<sub>0-144</sub>; log<sub>10</sub> cfu/mL·h)** |  |  |
| Stasis | 30.7% (0* - 40.9%) | 6.2 (1.8 - 13.3) |
| 1 log<sub>10</sub> reduction | 45.1% (33.7 - 53.6%) | 18.7 (7.1 - 31.4) |
| 2 log<sub>10</sub> reduction | 61% (51.7 - 72.9%) | 61.6 (36.6 - 95.9) |
| 3 log<sub>10</sub> reduction | 81.5% (67.8 - 91.1%) | 259.3 (154 - 440.2) |
| R<sup>2</sup> | 0.8386 | 0.8258 |

Exposures targets required for stasis, 1, 2 and 3 log<sub>10</sub> kill or total reduction in the bacterial response (area-under-bacterial-kill-curve, AUBKC<sub>0-144</sub>), applying MIC measurements using EUCAST methodology (static 2mg/L concentration of clavulanic acid). ~, No lower limit of confidence interval able to be interpolated.

**Table S6.**
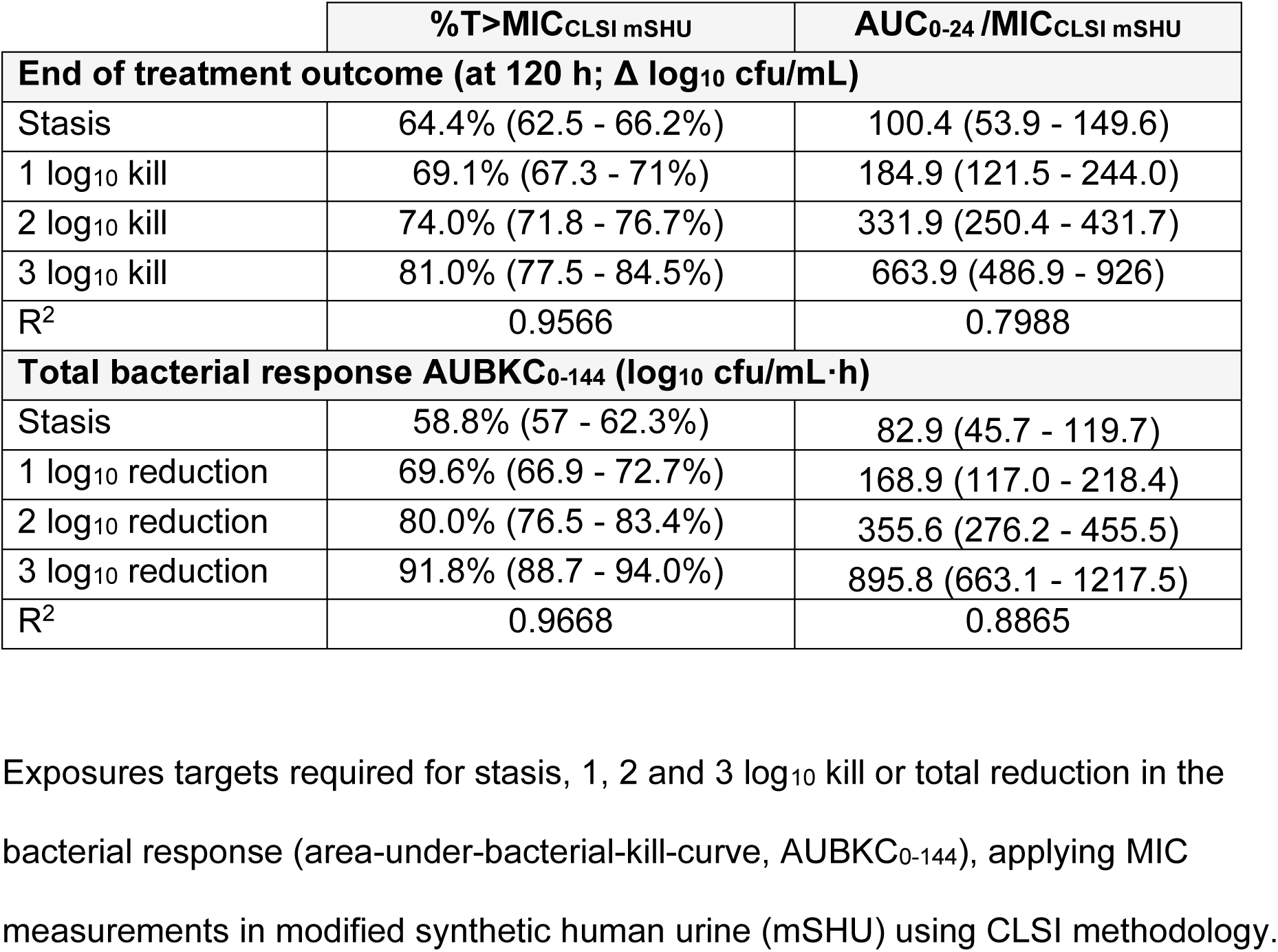
Amoxicillin-clavulanic acid urinary PK/PD targets with MICs determined in modified synthetic human urine.

|  | **%T>MIC<sub>CLSI mSHU</sub>** | **AUC<sub>0-24</sub> /MIC<sub>CLSI mSHU</sub>** |
|---|---|---|
| **End of treatment outcome (at 120 h; Δ log<sub>10</sub> cfu/mL)** |  |  |
| Stasis | 64.4% (62.5 - 66.2%) | 100.4 (53.9 - 149.6) |
| 1 log<sub>10</sub> kill | 69.1% (67.3 - 71%) | 184.9 (121.5 - 244.0) |
| 2 log<sub>10</sub> kill | 74.0% (71.8 - 76.7%) | 331.9 (250.4 - 431.7) |
| 3 log<sub>10</sub> kill | 81.0% (77.5 - 84.5%) | 663.9 (486.9 - 926) |
| R<sup>2</sup> | 0.9566 | 0.7988 |
| **Total bacterial response AUBKC<sub>0-144</sub> (log<sub>10</sub> cfu/mL·h)** |  |  |
| Stasis | 58.8% (57 - 62.3%) | 82.9 (45.7 - 119.7) |
| 1 log<sub>10</sub> reduction | 69.6% (66.9 - 72.7%) | 168.9 (117.0 - 218.4) |
| 2 log<sub>10</sub> reduction | 80.0% (76.5 - 83.4%) | 355.6 (276.2 - 455.5) |
| 3 log<sub>10</sub> reduction | 91.8% (88.7 - 94.0%) | 895.8 (663.1 - 1217.5) |
| R<sup>2</sup> | 0.9668 | 0.8865 |

## SUPPLEMENTARY FIGURES

**Figure S1.**
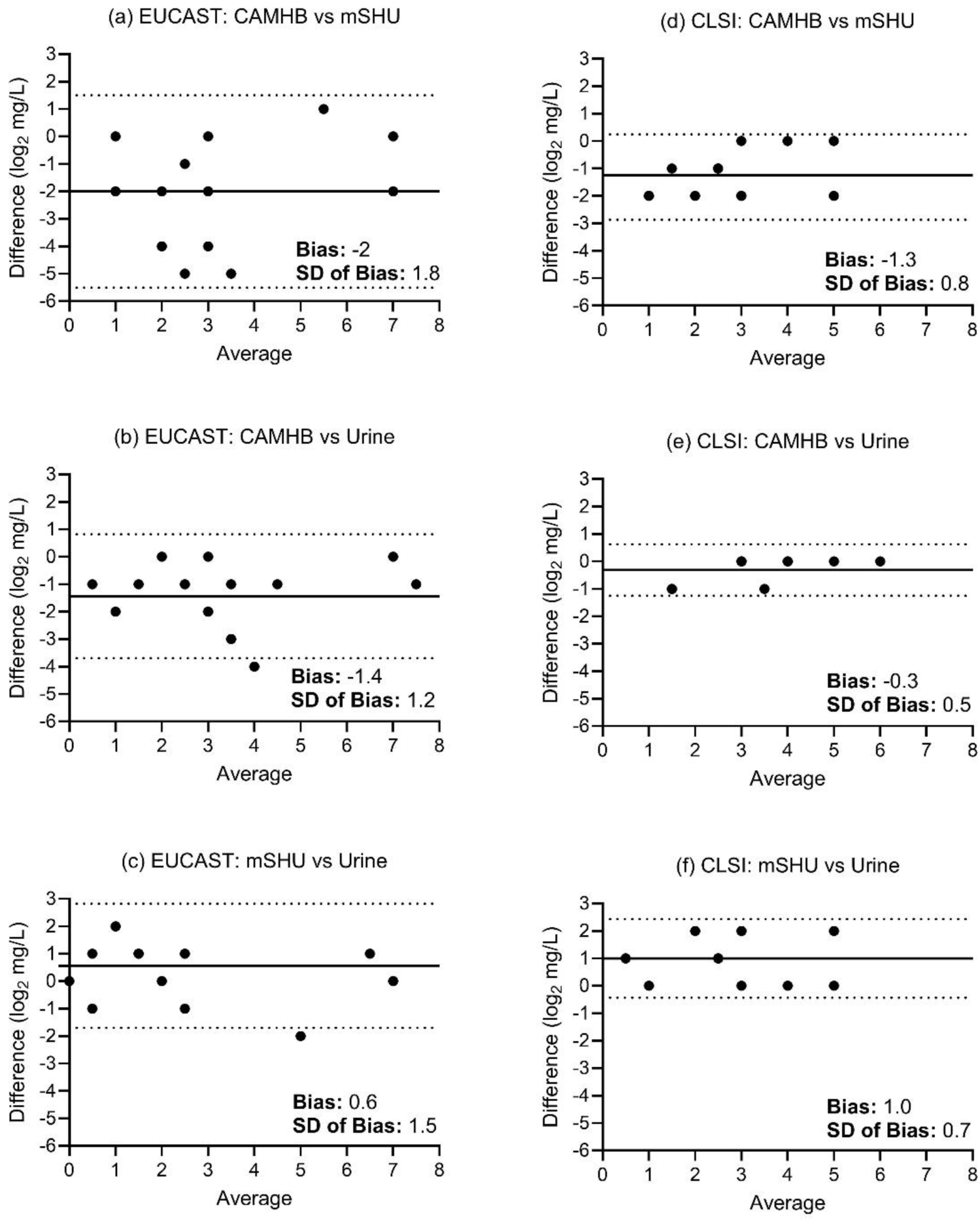
Bland-Altman analysis. Comparison between MICs in standard (CAMHB; Cation-adjusted Mueller-Hinton broth) and nonstandard (Pooled human urine; urine, modified synthetic human urine; mSHU) media. Bias and standard deviation of bias is reported.

**Figure S2.**
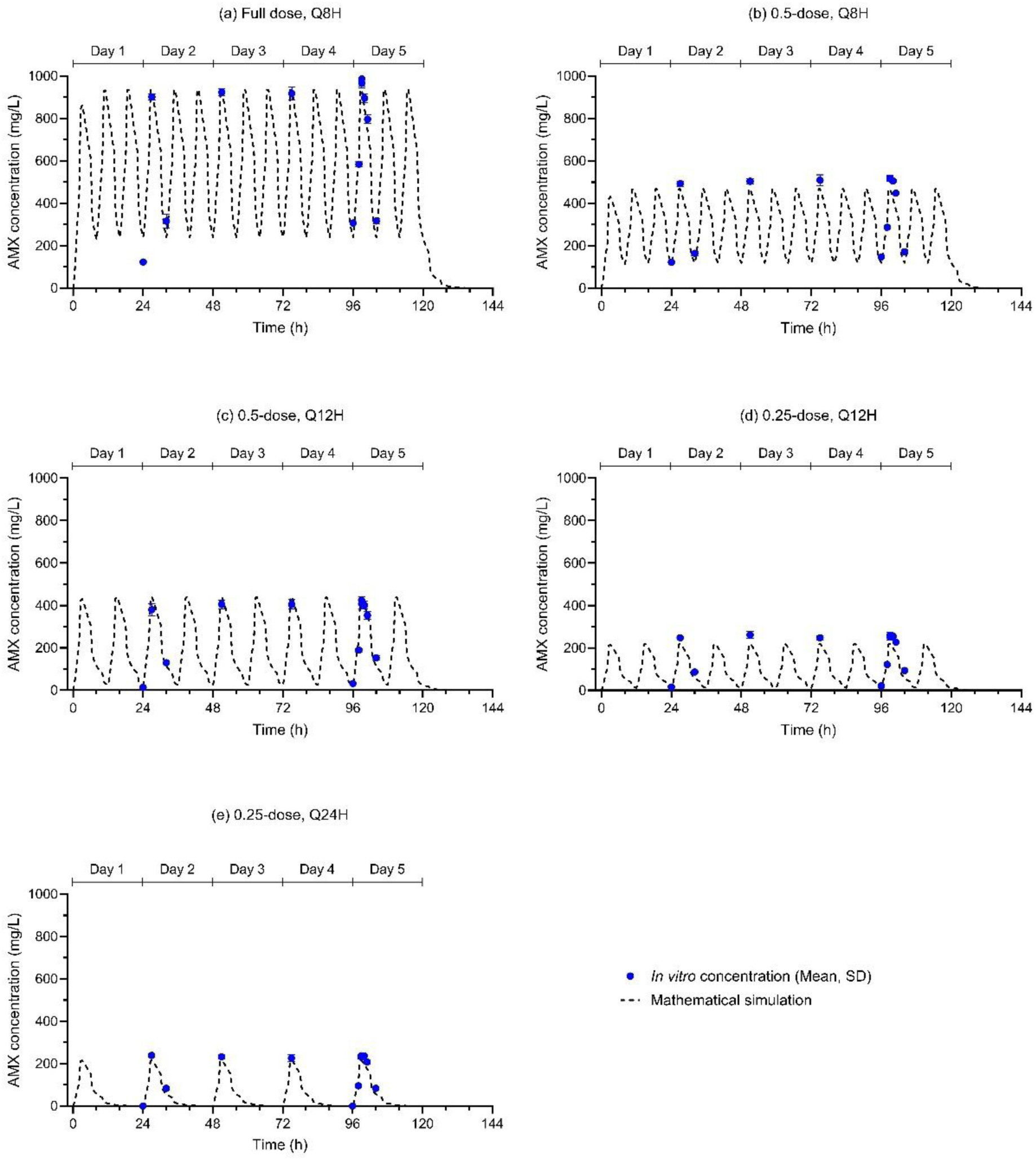
Amoxicillin exposures in the bladder infection model. The five simulated amoxicillin exposures in the bladder infection model alongside their measured concentrations. The five exposures were (a) standard exposure (8 hourly; C_max_ 935 mg/L; AUC_0-24_ 14272 mg/L), (b) intermediate exposure (8 hourly; C_max_ 467 mg/L; AUC_0-24_ 7073 mg/L), (c) low exposure (12 hourly; C_max_ 440 mg/L; AUC_0-24_ 4757 mg/L), (d) very low exposure (12 hourly; C_max_ 220 mg/L; AUC_0-24_ 2379 mg/L), and (d) minimal exposure (24 hourly; C_max_ 217 mg/L; AUC_0-24_ 1189 mg/L). Points indicate the measured *in vitro* amoxicillin concentrations. *In vitro* amoxicillin concentrations are reported as the mean and standard deviation of samples from three representative bladders, except for the C_max_ sample at 99 h in which all 16 bladders were sampled. AMX, Amoxicillin.

**Figure S3.**
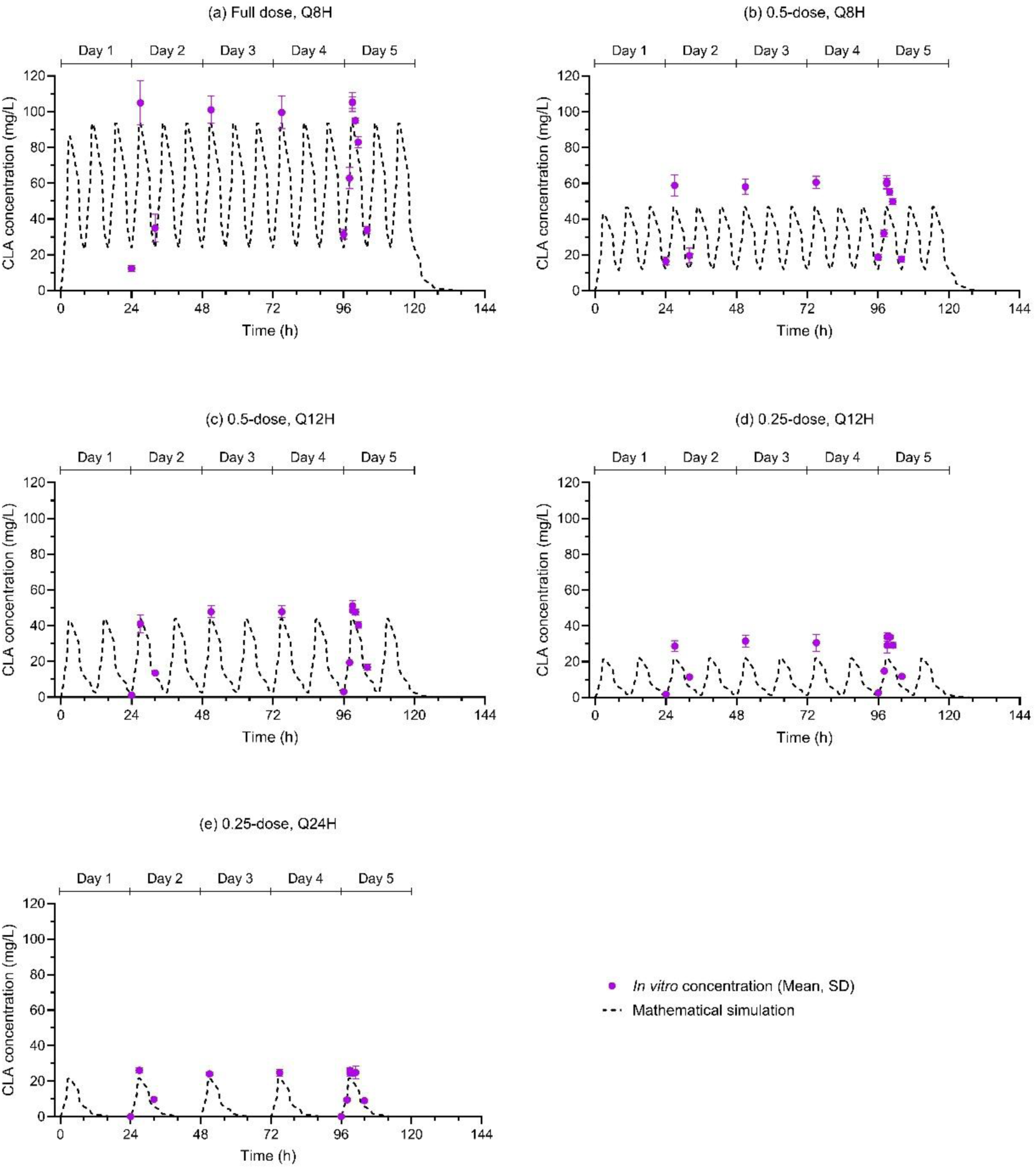
Clavulanic acid exposures in the bladder infection model. The five simulated clavulanic acid exposures in the bladder infection model alongside their measured concentrations. The five exposures were (a) standard exposure (C_max_ 93 mg/L; AUC_0-24_ 1427 mg/L), (b) intermediate exposure (C_max_ 47 mg/L; AUC_0-24_ 707 mg/L), (c) low exposure (C_max_ 44 mg/L; AUC_0-24_ 476 mg/L), (d) very low exposure (C_max_ 22 mg/L; AUC_0-24_ 238 mg/L), and (d) minimal exposure (C_max_ 22 mg/L; AUC_0-24_ 119 mg/L) Points indicate the measured *in vitro* amoxicillin concentrations. *In vitro* amoxicillin concentrations are reported as the mean and standard deviation of samples from three representative bladders, apart from the C_max_ sample at 99 h in which all 16 bladders were sampled. CLA, Clavulanic acid.

**Figure S4.**
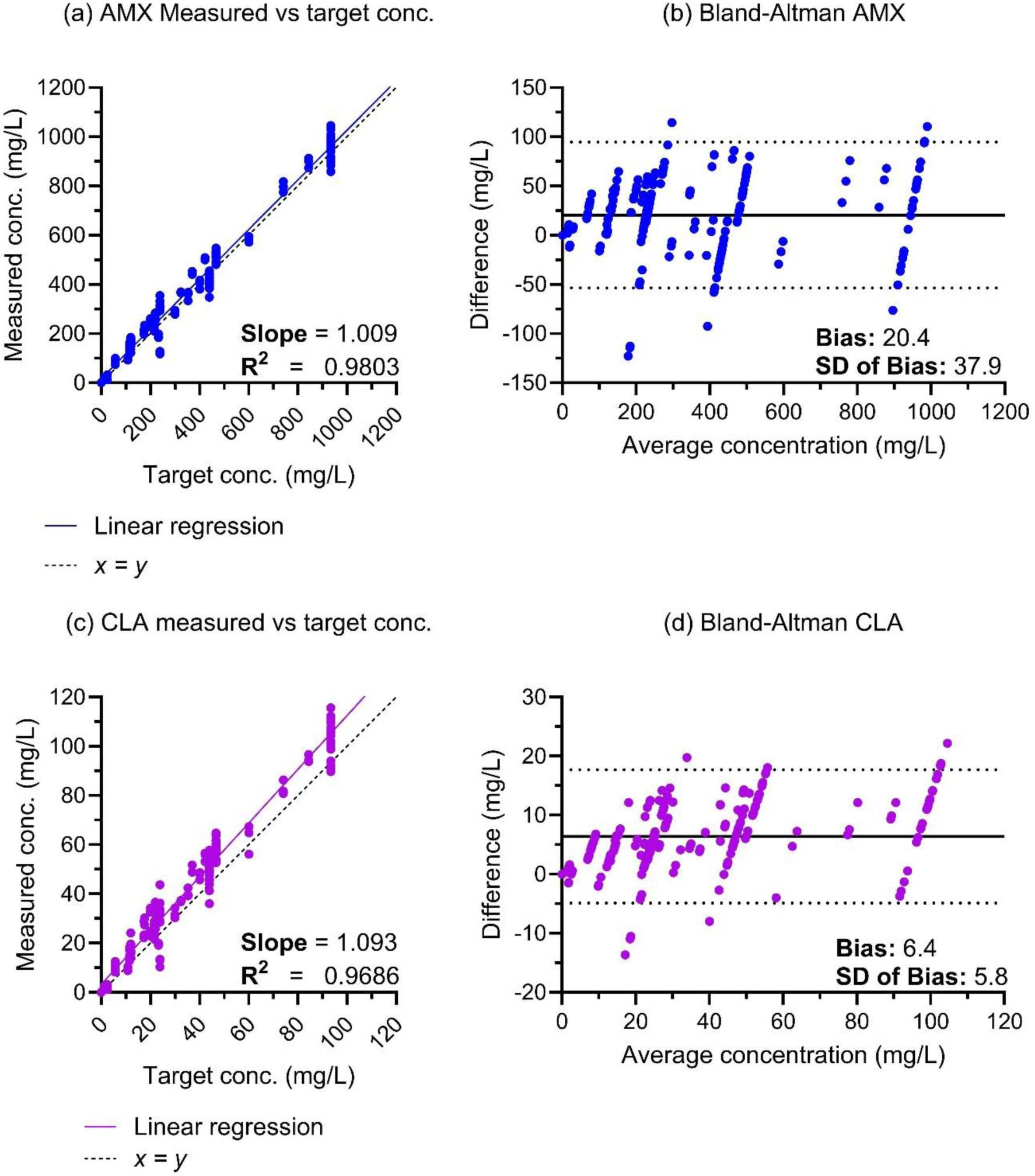
Measured versus target urinary amoxicillin and clavulanic acid concentrations. The accuracy of *in vitro* amoxicillin and clavulanic acid exposure in the bladder infection model assessed by (a) linear regression and (b) by Bland-Altman plot. Amoxicillin displayed a slope of 1.009 and R^2^ of 0.9803. Difference was calculated by measured amoxicillin concentration -simulated amoxicillin concentration and plotted against the average of the two. Bias for amoxicillin was 20.4mg/L (95% CI: -53.9 to 94.7). Accuracy of clavulanic dosing measured by (c) linear regression and (d) by Bland-Altman plot. Clavulanic acid displayed a slope of 1.093 and R^2^ of 0.9686. Bias for clavulanic acid was 6.4mg/L (95% CI: -4.9 to 17.7). AMX, Amoxicillin. CLA, Clavulanic acid.

**Figure S5.**
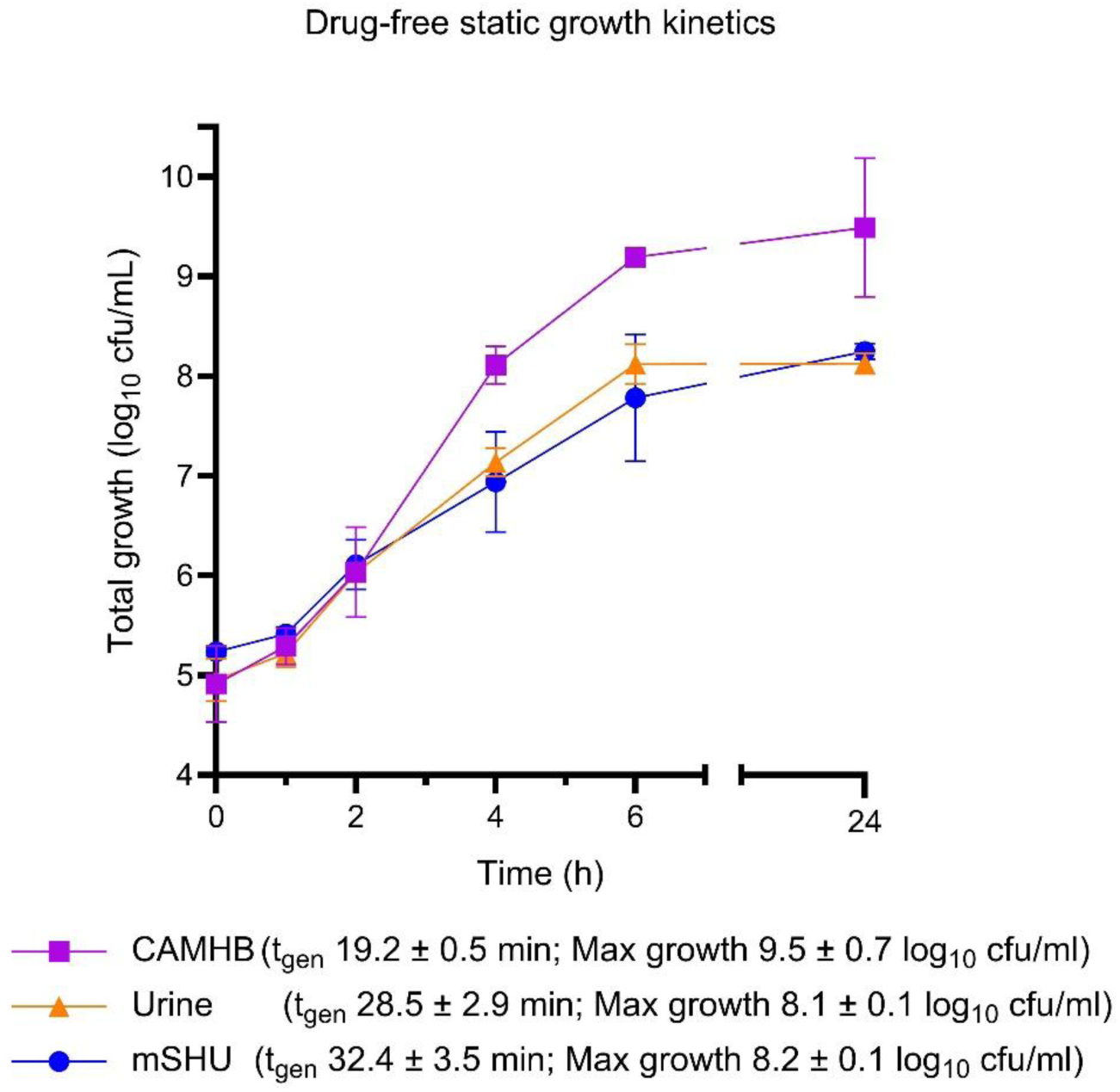
Drug-free static growth kinetics. 20mL of CAMHB, pooled human urine, and mSHU were inoculated with four isolates, ATCC 25922, ATCC 35218, CFT073 and UTI89 to a concentration of 10^5^ CFU/mL. Static growth kinetics of the four isolates were quantified over a 24 h period, with the mean and standard deviation of bacterial density of the four isolates presented above. Generation time and maximal growth capacity were reported.

**Figure S6.**
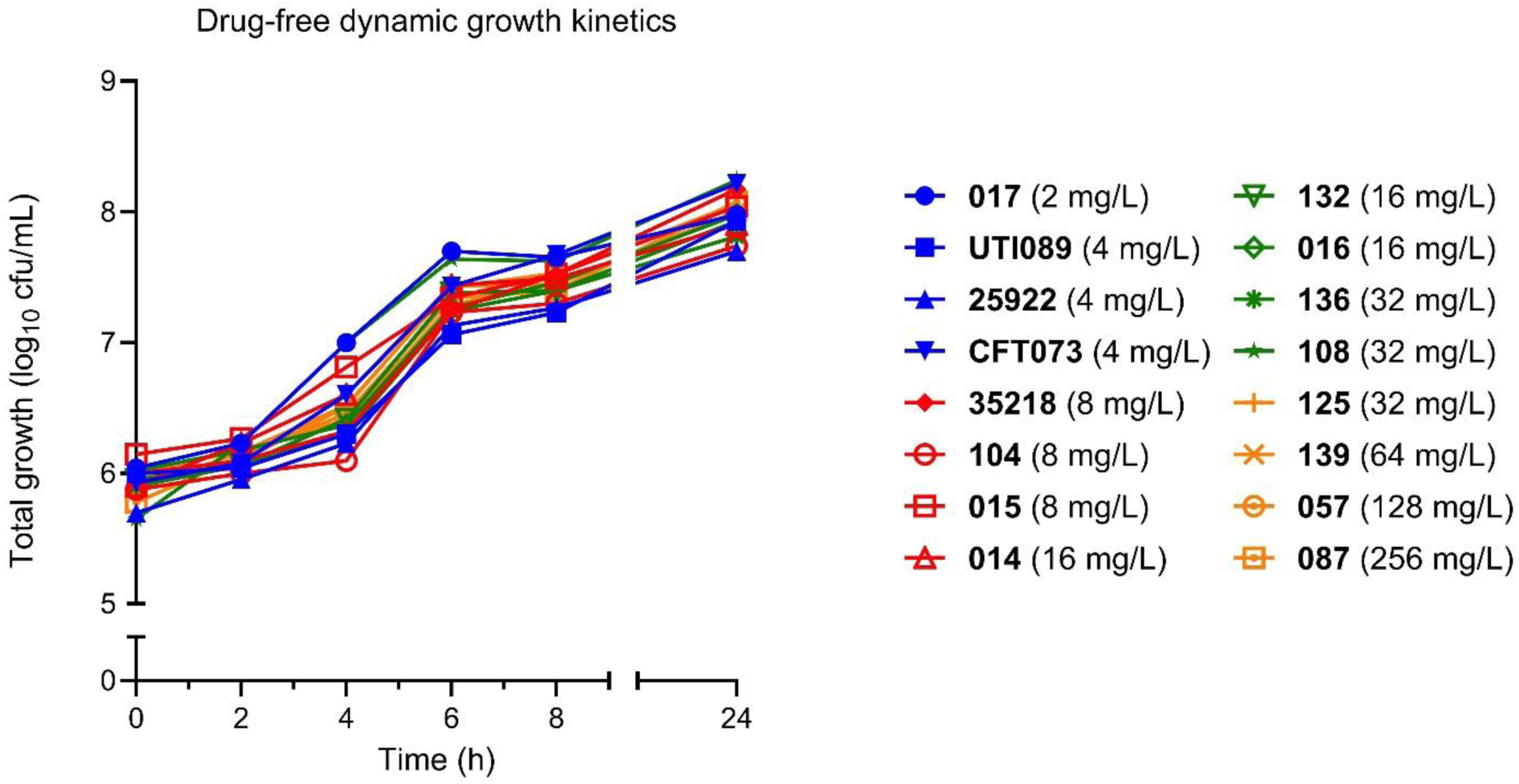
Drug free growth within the bladder infection model. After 24 hours of dynamic, drug free growth within the BIM, the 16 isolates displayed an average maximal growth capacity in the BIM of 8.0 ± 0.2 log_10_ cfu/mL.

**Figure S7.**
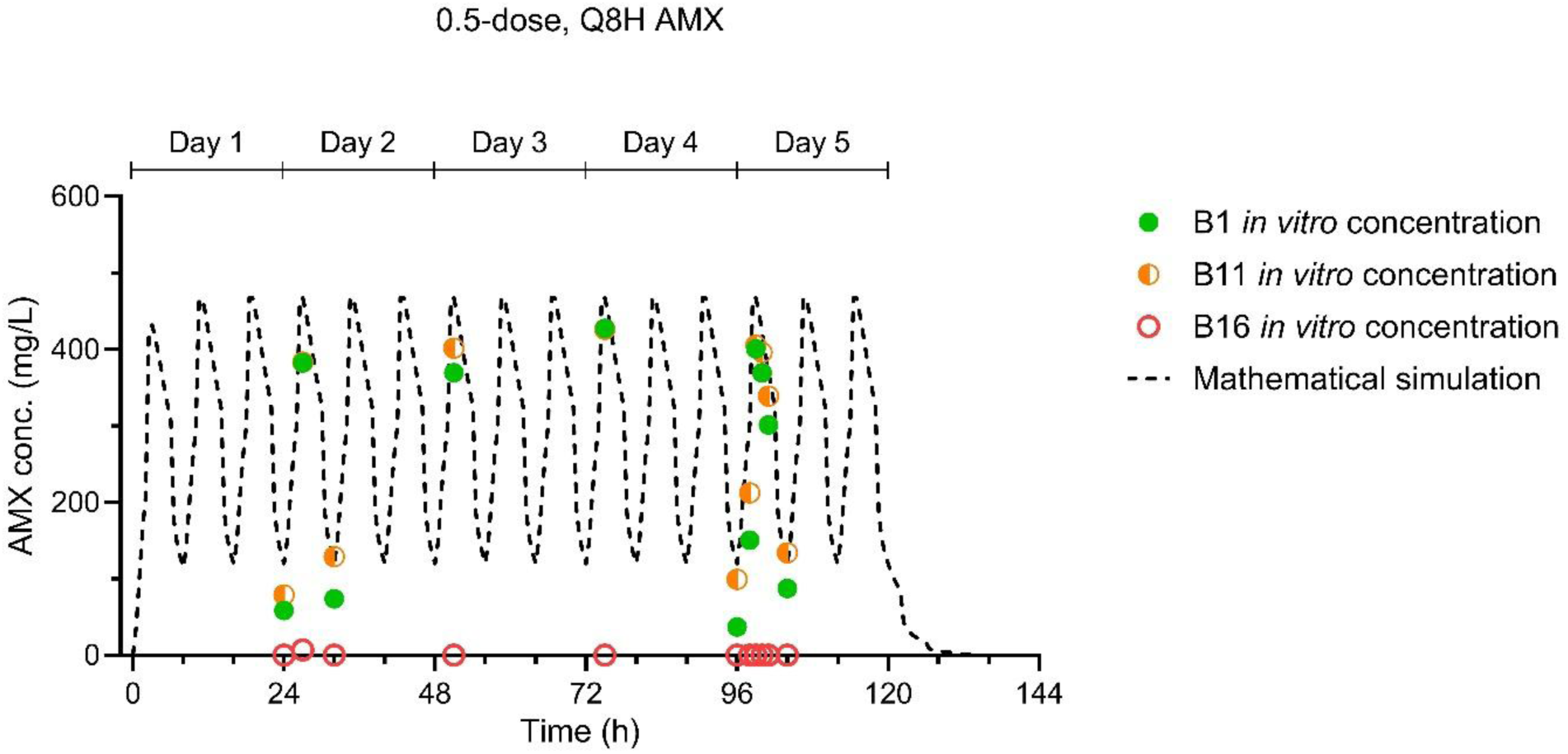
Amoxicillin concentrations during the amoxicillin-only exposure experiment. Measured *in vitro* concentrations as compared to the mathematical simulation for the amoxicillin only exposure. Simulated exposure matched the medium exposure with amoxicillin-clavulanic acid (8 hourly; C_max_ 467 mg/L; AUC_0-24_ 7073 mg/L). Each point represents a single sample from one of three representative bladders. Each antibiotic sample came from bladders 1, 11 and 15, which contained isolates ECO 017, ECO 136, and ECO 057 respectively. AMX, Amoxicillin.

**Figure S8.**
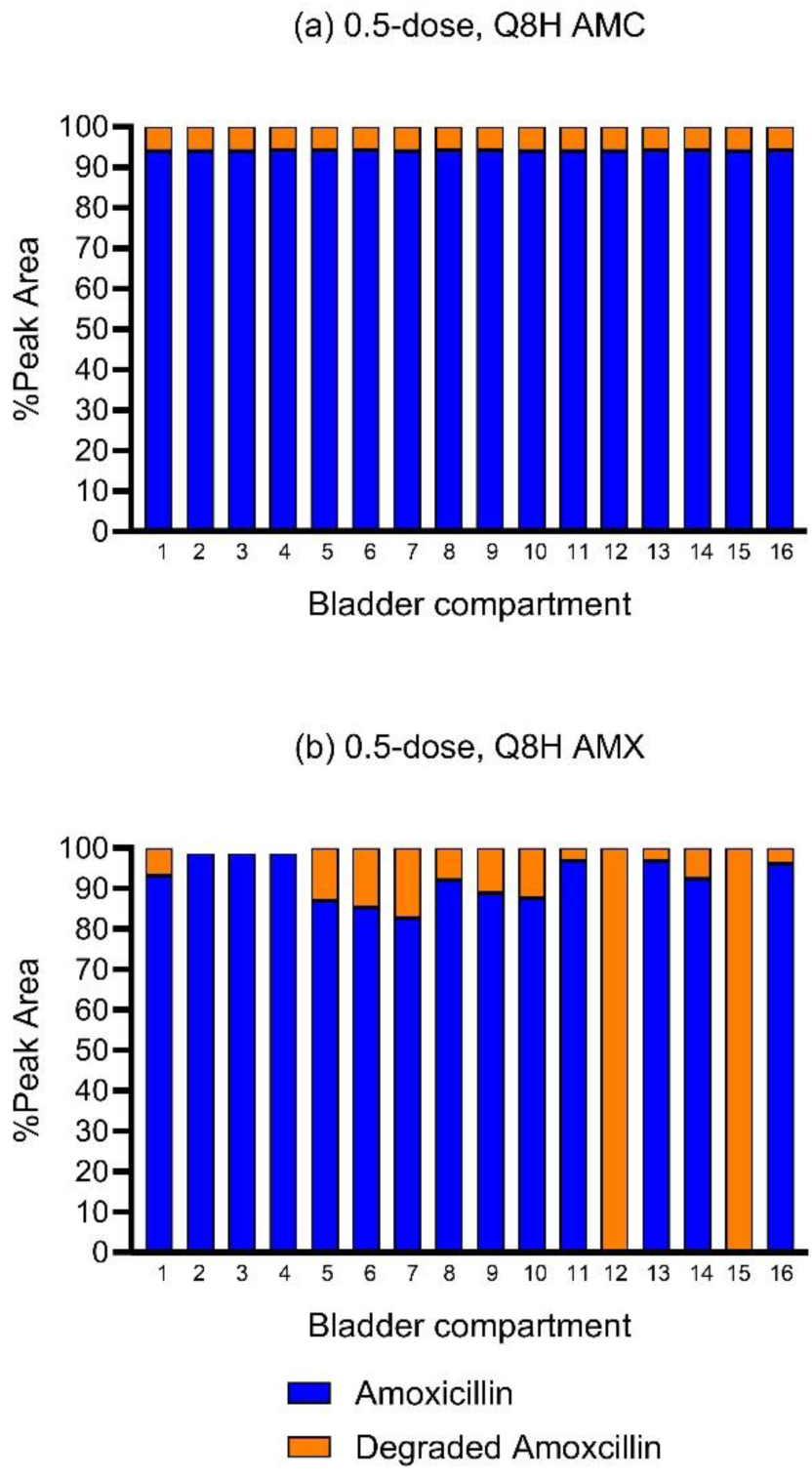
**HPLC-PDA peak analysis of amoxicillin and its degradation products**. AMC, Amoxicillin-clavulanic acid. AMX, Amoxicillin. Assessment of the day 5 peak concentration measurement from all 16 bladder compartments following the (a) half-dose 8-hourly amoxicillin-clavulanic acid experiment and (b) half-dose 8-hourly amoxicillin-only experiment. The amoxicillin degradation peaked at 1.6 min. There was an additional small peak at approximately 1.7 minutes that was not classified as a degradation peak; therefore, the area associated with this peak accounts for the difference when the amoxicillin and degradation peak percentages do not sum to 100%.

**Figure S9.**
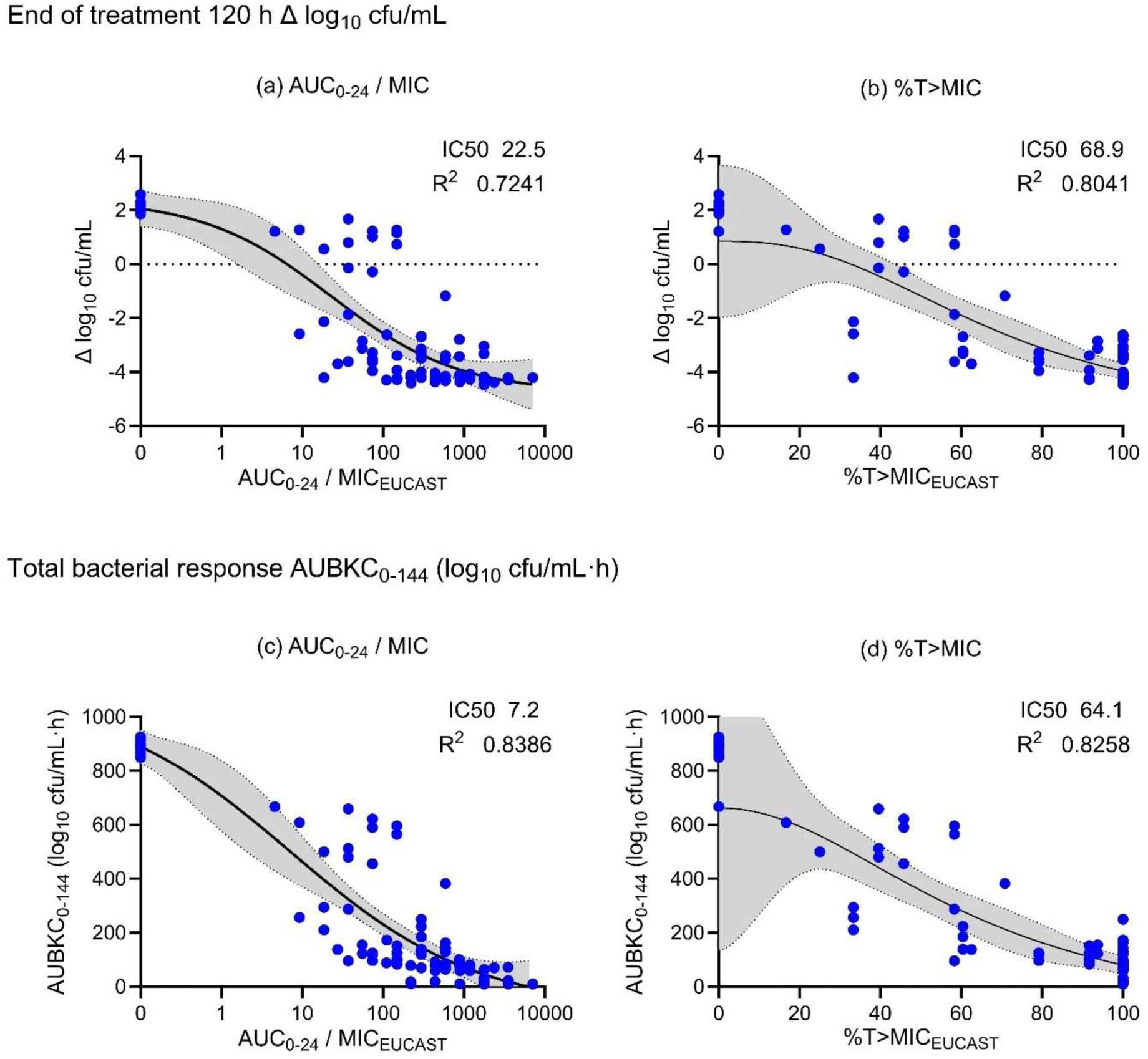
Exposure response relationships for amoxicillin-clavulanic acid utilizing MICs measured by EUCAST methodology (fixed 2 mg/L clavulanic acid). The four-way parametric E_max_ models above detail the primary (end of treatment Δ log_10_ cfu/mL) and secondary outcome (AUBKC_0–144_). Graphs have not been constrained. Shaded area indicates the 95% confidence band. The 50% maximal inhibitory concentration (IC_50_) and coefficient of determination (R^2^) are reported.

**Figure S10.**
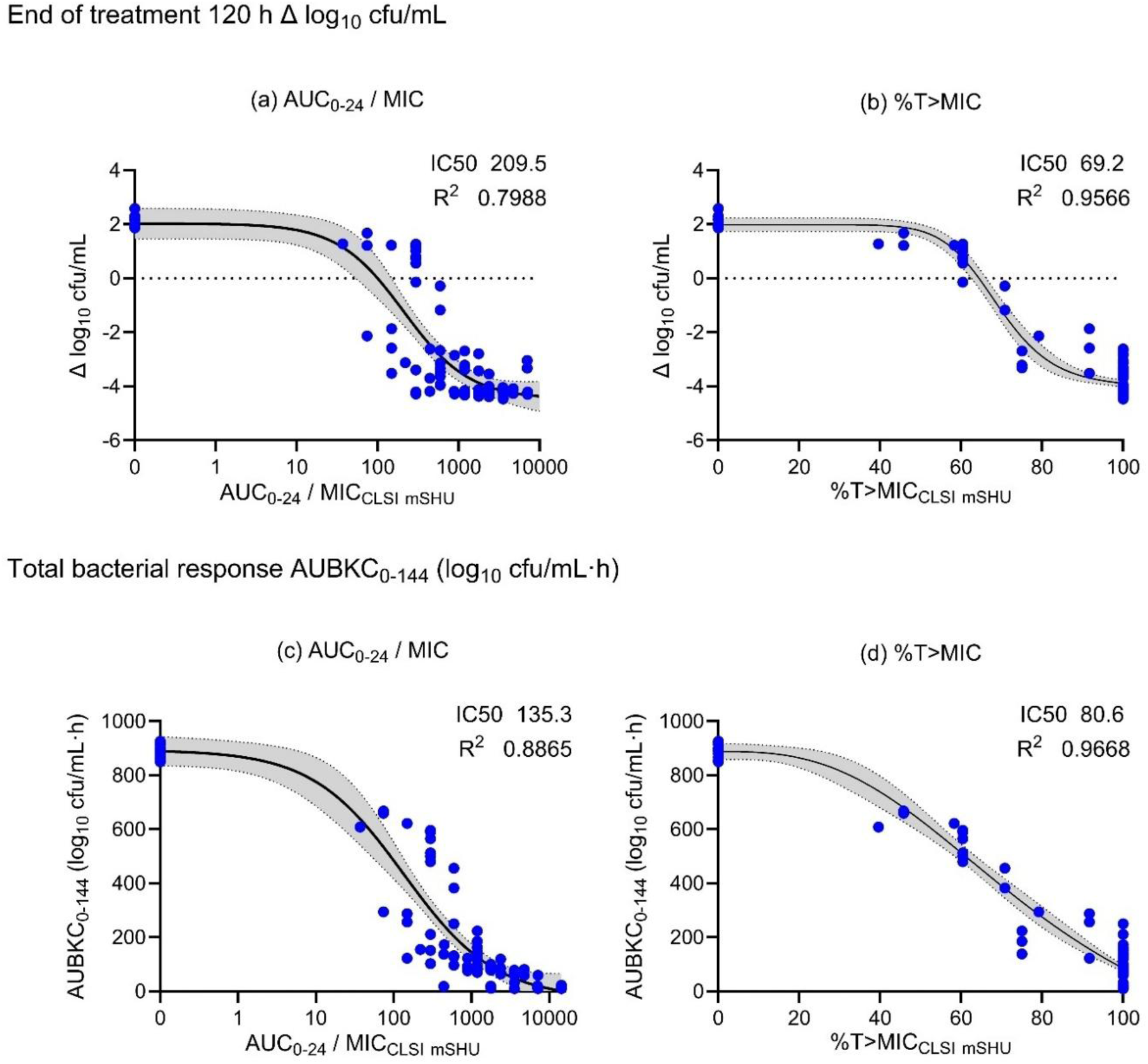
Exposure response relationships for amoxicillin-clavulanic acid utilizing MICs tested in mSHU and measured by CLSI method (2:1 ratio of amoxicillin to clavulanic acid). The four-way parametric E_max_ models above detail the primary (end of treatment Δlog_10_ CFU/mL) and secondary outcome (AUBKC_0–144_). Graphs have not been constrained. Shaded area indicates the 95% confidence band. The 50% maximal inhibitory concentration (IC_50_) and coefficient of determination (R^2^) are reported.

**Figure S11:**
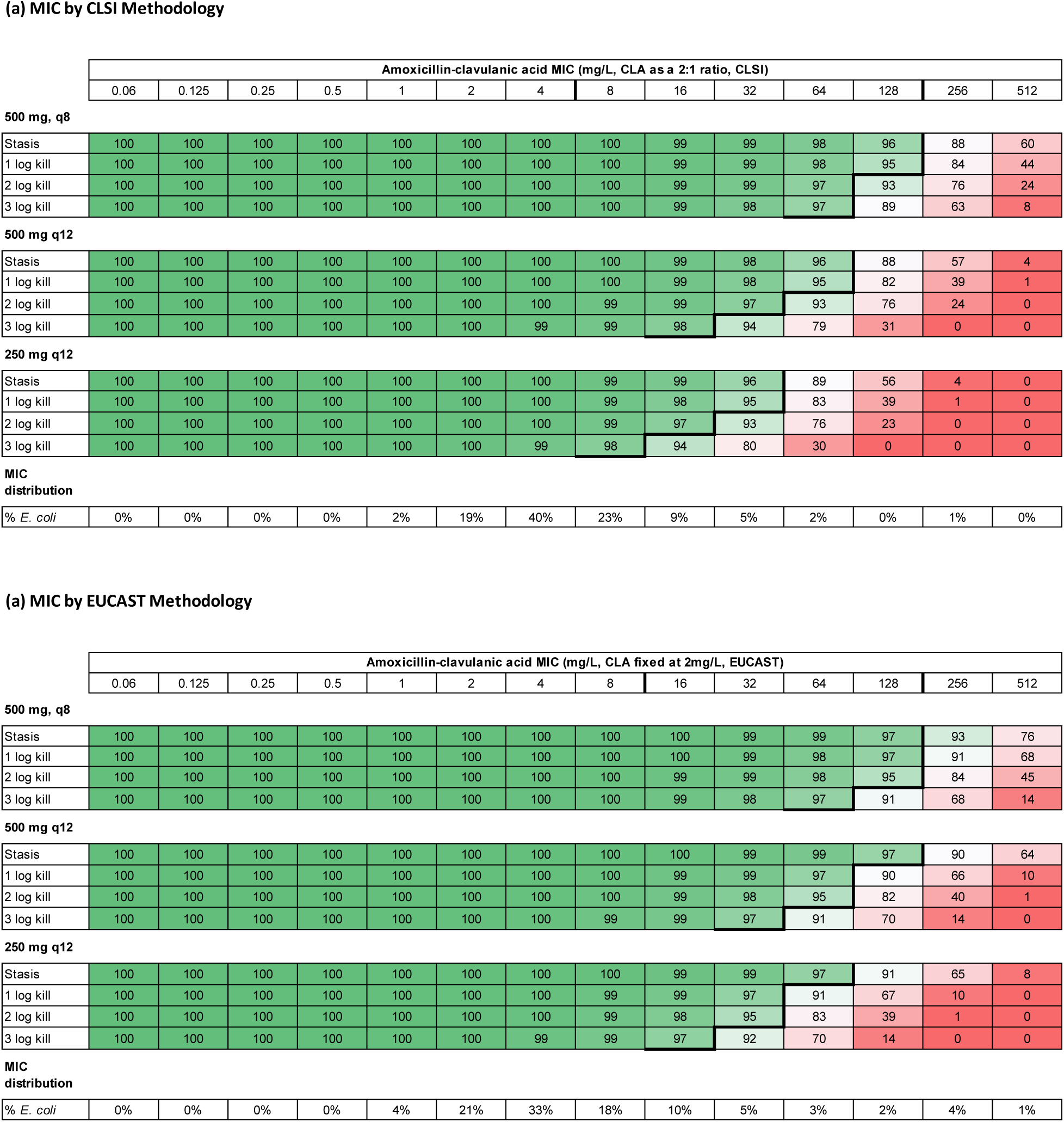
Monte Carlo Simulations. Probability of target attainment in 5000 virtual patients for standard 8-hourly dosing and following reduced dose and dosing frequency (± 50% variation in exposure). Applied %T_>MIC_ targets determined for the end of treatment bacterial responses (stasis and log kill), using MIC values determined by (a) CLSI methodology and (b) EUCAST methodology. This data highlights the MIC values of isolates in which >95% of virtual patients are expected to achieve the end of treatment PK/PD target.

